# High-resolution cryo-EM structure of urease from the pathogen *Yersinia enterocolitica*

**DOI:** 10.1101/2020.04.28.065599

**Authors:** Ricardo D. Righetto, Leonie Anton, Ricardo Adaixo, Roman P. Jakob, Jasenko Zivanov, Mohamed-Ali Mahi, Philippe Ringler, Torsten Schwede, Timm Maier, Henning Stahlberg

## Abstract

Urease converts urea into ammonia and carbon dioxide and makes urea available as a nitrogen source for all forms of life except animals. In human bacterial pathogens, ureases also aid in the invasion of acidic environments such as the stomach by raising the surrounding pH. Here, we report the structure of urease from the pathogen *Yersinia enterocolitica* at better than 2 Å resolution from cryo-electron microscopy. *Y. enterocolitica* urease is a dodecameric assembly of a trimer of three protein chains, ureA, ureB and ureC. The high data quality enables detailed visualization of the urease bimetal active site and of the impact of radiation damage. Our data are of sufficient quality to support drug development efforts.

## Introduction

Ureases are nickel-metalloenzymes produced in plants, fungi and bacteria, but not in animals. They facilitate nitrogen fixation by metabolizing urea, but in some pathogenic bacteria they serve a dual function and consume ammonia to promote survival in acidic environments. This function is vital during host infection where the bacteria have to survive the low pH of the stomach (Gripenberg-Lerche *et al*., 2000), and of phagosomes in host cells (Young *et al*., 1996; Maroney & Ciurli, 2014). The relevance of ureases in early stages of infection renders them attractive targets for novel anti-microbials.

Ureases catalyze the breakdown of urea into ammonia and carbamate at a rate 10^14^ to 10^15^ times faster than the non-catalyzed reaction **(Fig. 1a)**, and are arguably the most efficient hydrolases (Maroney & Ciurli, 2014). In a second non-catalyzed step, carbamate is spontaneously hydrolyzed to yield another molecule of ammonia as well as one molecule of bicarbonate (Mazzei *et al*., 2017). Jack bean urease was the first enzyme to be crystallized, offering evidence that enzymes are proteins (Sumner, 1926) and was the first metalloenzyme to be shown to use nickel in its active site (Dixon *et al*., 1975). All ureases characterized to date share the architecture of their active site (Kappaun *et al*., 2018). Two Ni^2+^ ions in the active site are coordinated by a carbamylated lysine, four histidines and one aspartate. Urea first interacts with Ni(1) through its carbonyl oxygen, making urea more available for nucleophilic attacks. The amino nitrogen then binds to Ni(2) and a proton is transferred from a water molecule to the amine in a nucleophilic attack on the carbonyl carbon of urea. The active site is closed off during the reaction by a conformationally variable helix-turn-helix motif, referred to as the mobile flap. Despite their common catalytic mechanism, ureases display different chain topologies and higher order oligomeric assemblies **(Fig. 1b,c)**. Jack bean urease assembles into an oligomer of a single type of polypeptide chain with D3 symmetry ([[α]_3_]_2_ D3) **(Fig. 1b,c)** (Maroney & Ciurli, 2014). The urease of the human pathogen *Helicobacter pylori* is composed of two types of polypeptide chains (ureA, ureB) and assembles into a dodecamer (tetramer-of-trimers) with tetrahedral symmetry ([[αβ]_3_]_4_ T) **(Fig. 1 b,c)**. The ability of ureases to raise the pH of their environment benefits *H. pylori* in colonizing the stomach and downstream gut, causing gastric ulcers (Kappaun *et al*., 2018). The urease of the opportunistic pathogen *Klebsiella aerogenes* (Podschun & Ullmann, 1998) assembles into a heterotrimer of three proteins ureA, ureB and ureC, which in turn oligomerizes into a trimer ([αβγ]_3_ C3) (Jabri *et al*., 1995) **(Fig 1b,c)**. This oligomeric assembly is common also to most other structurally characterized bacterial ureases (Maroney & Ciurli, 2014).

**Figure 1.**
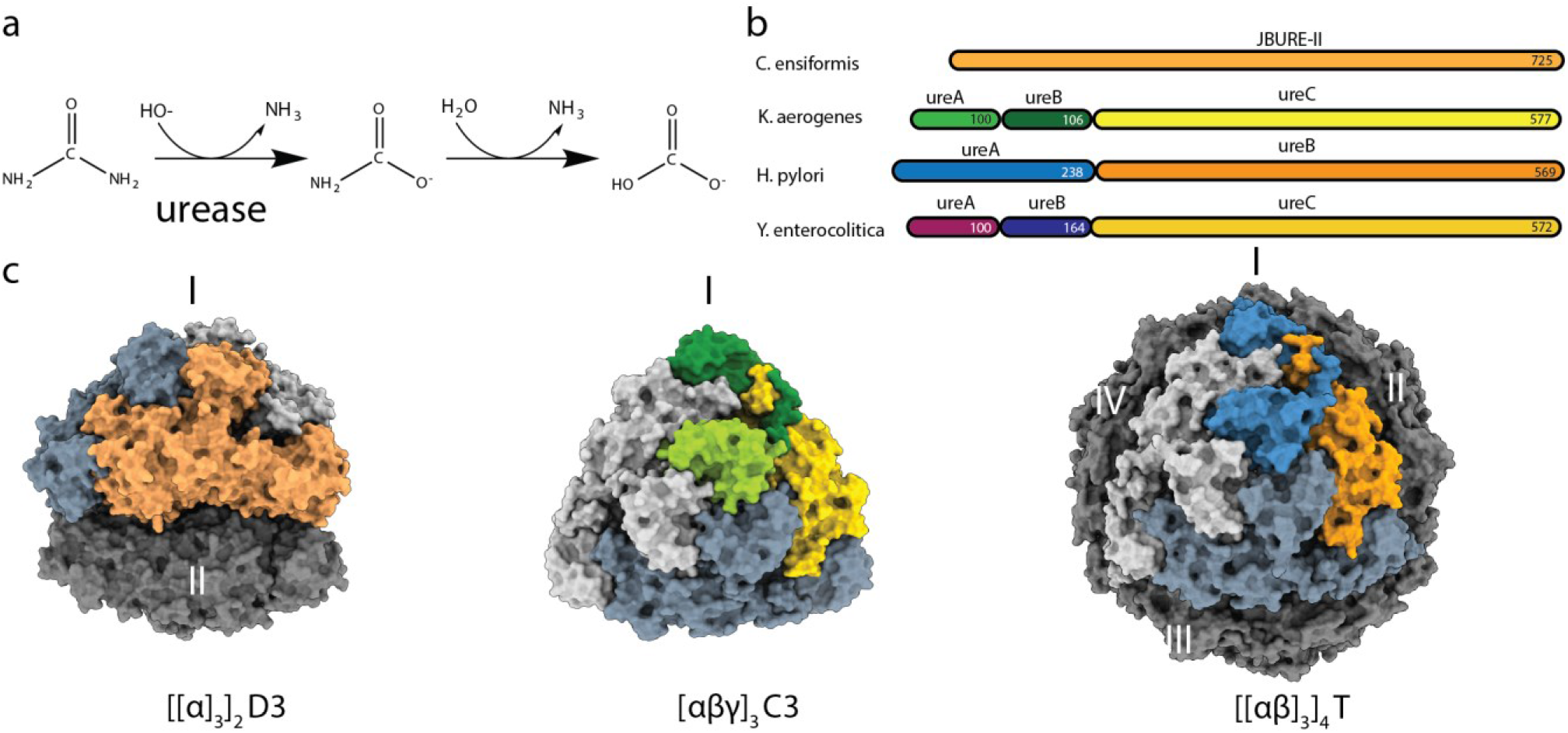
Protein architectures and oligomeric assemblies of ureases. **a)** Schematic of biochemical reaction catalyzed by urease. **b)** Protein architecture of urease functional unit of *C. ensiformis* (Jack bean urease), *K. aerogenes, H. pylori* and *Y. enterocolitica*. **c)** Surface representation of oligomeric assembly of urease in *C. ensiformis* (PDB: 3LA4), *K. aerogenes* (PDB: 1EJW), *H. pylori* (PDB: 1E9Z), respectively. Proteins are color-coded as in **b)**. Oligomeric state of urease assembly is indicated at the bottom and the trimeric assemblies are indicated in roman numerals.

*Yersinia enterocolitica* is the causative agent of yersiniosis, a gastrointestinal infection, reactive arthritis and erythema nodosum. The infection spreads to humans through consumption of contaminated food with pigs being one of the largest reservoirs of *Y. enterocolitica*. The symptoms of yersiniosis are fever, abdominal pain, diarrhea and/or vomiting and is one of the most reported enteritis in some countries, although outbreaks are rare (Drummond *et al*., 2012). *Y. enterocolitica* is a facultative intracellular bacterium and can survive in very different environments. In the presence of urea *Y. enterocolitica* can tolerate extremely acidic conditions (Young *et al*., 1996). *Y. enterocolitica* urease comprises three polypeptide chains (ureA, ureB and ureC), an architecture similar to that of *K. aerogenes* urease **(Fig. 1b)**.

Here we present the structure of *Y. enterocolitica* urease at an overall resolution of 1.98 Å, which was achieved using recent advances in cryo-EM data collection and processing. The structure shows that *Y. enterocolitica* urease assembles into a dodecameric hollow sphere with [[αβγ]_3_]_4_ T oligomeric assembly structure **(Fig. 2)**. A tightly embedded kinked loop is interacting with neighboring domains and is potentially responsible for the assembly of the oligomer. The data allows model building of the active site carbamylated lysine, visualization of radiation damage to the nickel-metal center, and hydration networks throughout the protein. Our work highlights the potential of cryo-EM for structure-based drug discovery, specifically for large enzyme assemblies, which have been historically difficult to analyze by X-ray crystallography.

**Figure 2.**
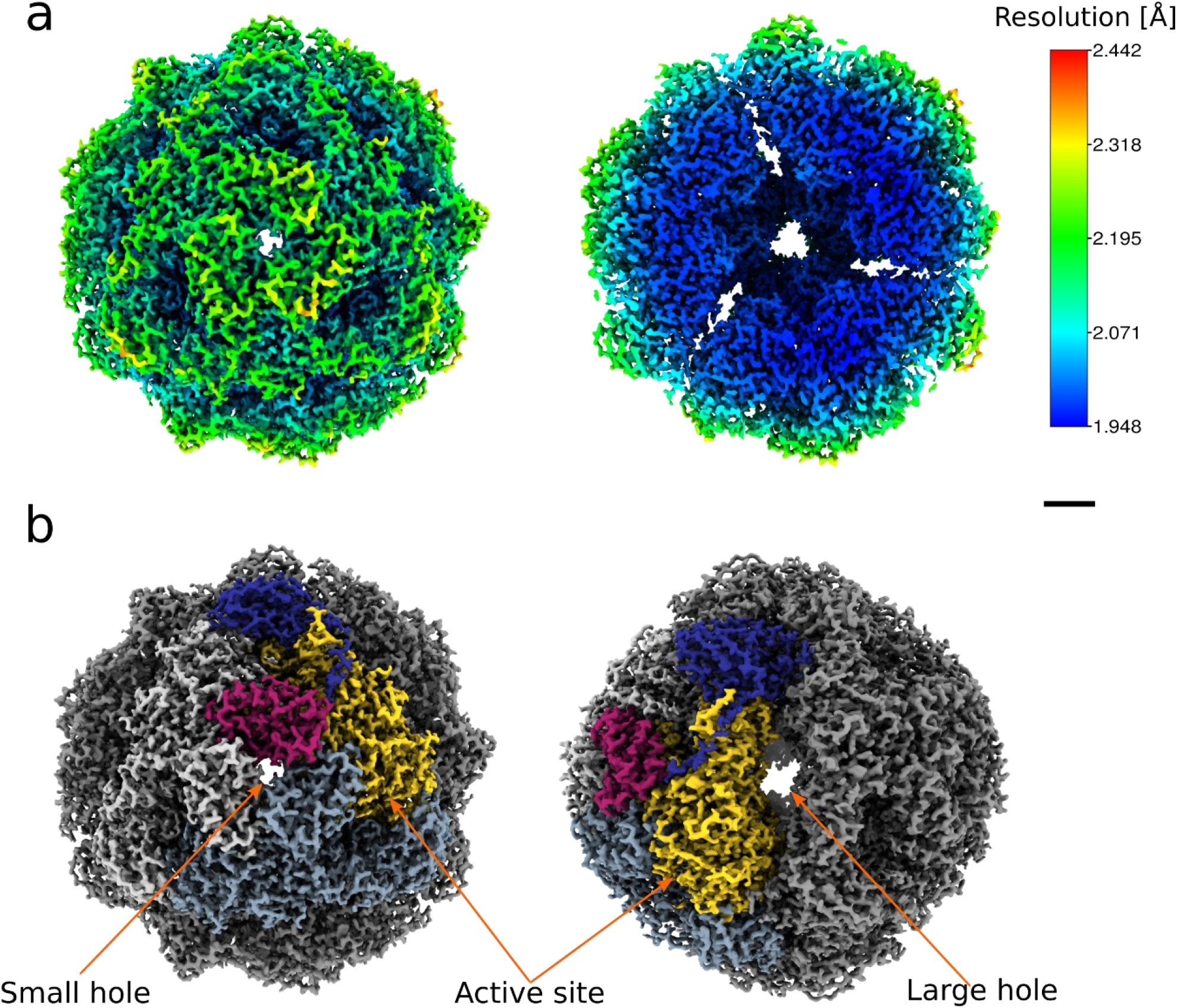
Cryo-EM analysis of the *Y. enterocolitica* dodecameric urease assembly. **a)** The cryo-EM map filtered and colored by local resolution (left), and a slice cut through the map to show the internal details (right). **b)** The assembly architecture highlighted on the map. The three chains that form the basic hetero-trimer are shown in different colors, with the other hetero-trimers shown in shades of gray. Two different views are shown to indicate the location of the small and larger holes at the interfaces, as well as the active site. Scale bars: 20 Å.

## Results & Discussion

### Structure determination of *Y. enterocolitica* urease by cryo-EM

We have used single particle cryo-EM to determine the structure of the fully assembled *Y. enterocolitica* urease. We acquired 4,494 movies of urease particles using a Titan Krios transmission electron microscope (TEM) equipped with a K2 direct electron detector and an energy filter (see **Methods** for details). Approximately half of the movies (2,243) were acquired by illuminating three locations (“shots”) per grid hole using beam-image shift in order to speed up the data collection (Cheng *et al*., 2018), whereas the remaining movies were recorded without this feature i.e. just a single shot at the center of the hole. This allowed us to measure and assess the extent of beam tilt and other optical aberrations, as well as the behavior of sample drift between each condition and beam-image shift position. Typical micrographs from the imaged grids are shown in **Supp. Fig. 1a** and a summary of data collection information is given in **Supp. Tab. 1**.

Each dataset was processed separately for 3D reconstruction following the strategy depicted in **Supp. Fig. 2**. The first obtained 3D map, at an overall resolution of 2.6 Å, revealed that this urease assembly is a dodecamer of tetrahedral (T) symmetry with a diameter of approximately 170 Å. The separate processing of each dataset yielded refined 3D maps at nominal resolutions of 2.10 Å and 2.20 Å for the multi-shot and single-shot cases, respectively (see **Methods**).

For comparison, we also processed the merged set of particles from both datasets altogether. We observed on the 2D class averages a preferential orientation for the three-fold symmetric view of urease, and also the presence of isolated monomers and broken assemblies **(Supp. Fig. 1b)**. The presence of such incomplete assemblies was further confirmed by performing 3D classification without imposing symmetry, as shown in **Supp. Fig. 1c**. The 3D class corresponding to the complete dodecameric assembly of urease contained 119,020 particles, of which 69,512 (58.4%) came from the multi-shot and 49,518 (41.6%) from the single-shot dataset. With respect to the number of particles picked from each dataset, 64.7% of the particles from the multi-shot and 56.8% from the single-shot datasets were retained at this stage and throughout the final reconstruction. While coma-free alignment was performed and active beam-tilt compensation in SerialEM (Schorb *et al*., 2019) was used on our data collections, after performing beam tilt refinement in RELION-3 (Zivanov *et al*., 2018) we observed that the single-shot case has a residual beam-tilt higher than the smallest residual observed in the multi-shot case **(Supp. Tab. 2)**. These two values are however very close to zero and are possibly within the error margin of the *post hoc* beam tilt refinement procedure.

The reduced need to move the specimen stage in beam-image shift mode not only speeds up data collection but also minimizes stage drift. The second and third shots from the multi-shot dataset have comparatively less drift than both the first multi-shot and the single shot, as suggested by the parameter values obtained from the Bayesian polishing training (Zivanov *et al*., 2019a) on each beam-tilt class separately **(Supp. Tab. 3)**. As all the three multi-shots are taken in nearby areas within the same foil hole, this observation is consistent with the annealing of the vitreous ice layer and its carbon support after pre-irradiating the specimen as reported previously (Brilot *et al*., 2012).

At this point, the nominal resolution of the map after 3D refinement was 2.05 Å. Finally, correcting for residual higher-order aberrations in CTF refinement (Zivanov *et al*., 2019b) **(Supp. Fig. 3)** yielded a map at a global resolution of 1.98 Å **(Supp. Fig. 4a)**. Local resolution estimation reveals that the core of the map is indeed at this resolution level or better **(Fig. 2a and Supp. Fig. 4b)**, and the local resolution-filtered map was then used for model building as explained in the next section. Despite the twelve-fold symmetry of the urease assembly, a limiting factor in the resolution of the map is the strong presence of preferential orientation, as confirmed by the plot of the final orientation assignments (**Supp. Fig. 4c**). The estimated angular distribution efficiency is 0.60 (Naydenova & Russo, 2017). An overview of the cryo-EM map and its main features are depicted in **Supp. Mov. 1.**

### *Y. enterocolitica* urease assembles as a tetramer of trimers

Model building was initiated from available crystallographic models with subsequent fitting and refinement against the cryo-EM map. The model was built and refined for one asymmetric unit containing one copy of the ureA, ureB and ureC protein each. The model was then expanded using NCS (see **Methods**). The complete model covering the whole oligomeric assembly contains 9,552 residues, 3,672 waters and 24 nickel ions (two per active site, twelve active sites) **(Table 1)**. The quality of the model was assessed with the cryo-EM validation tools in the PHENIX package (Afonine, Klaholz *et al*., 2018). The map allowed for the building of all residues of ureA (1-100), and residues 31-162 of ureB and 2-327/335-572 of ureC **(Supp. Fig. 5)**. The hetero-trimer formed by the three protein chains (ureA, ureB, ureC) **(Fig. 1b)** oligomerizes into a homo-trimer. The homo-trimer is arranged in a tetramer-of-trimers making the full complex a dodecamer of the hetero-trimer **(Fig. 2b)**. There are four large oval shaped holes between the trimers (64 Å long, 12 Å wide, high electrostatic potential) and four smaller holes at the center of the trimer with a diameter of 6 Å (low electrostatic potential), as shown in **Fig. 2b**, and the center of the enzyme assembly is hollow **(Fig. 2).** The holes provide ample opportunity for diffusion of the uncharged substrate and product, and the hollow inside potentially leads to local increase of reaction product. The assembly has the same symmetry as the urease homologue in *H. pylori*, which was postulated to increase stability and/or resistance to acidic environments (Ha *et al*., 2001).

**Table 1.**
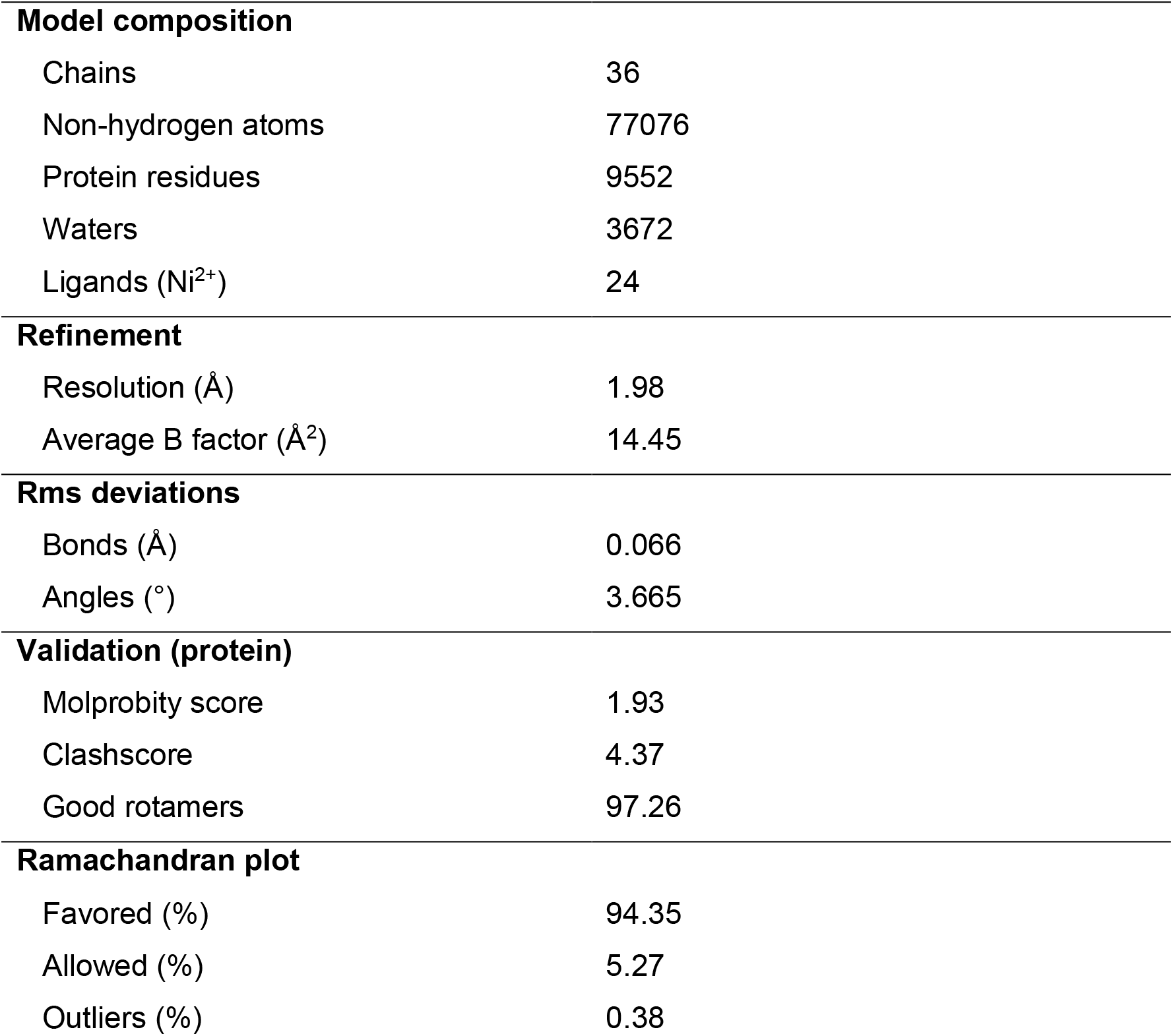
Model building and refinement. Statistics shown for full assembly calculated from the asymmetric unit using NCS.

For analysis of the protein sequences, the ConSurf Server (Berezin *et al*., 2004; Ashkenazy *et al*., 2016) was used with the sample list of homologs option to get a diverse set of 150 sequences. The protein chains of *Y. enterocolitica* urease are highly conserved across different organisms. The ureA chain is split after a LVTXXXP motif and is 99-100 amino acids long in most cases, with a sequence identity of 55.7%. The ureB chain of *Y. enterocolitica* has between 20 and 30 N-terminal amino acids more compared to the other sequences (except *Kaistia sp*. SCN 65-12), which share an identity of 51.5%. This N-terminal extension is located on the outside of the holoenzyme where ureA and ureB chain split occurs and are too disordered to be modeled in the structure **(Supp. Fig. 5)**. The charges and properties of this stretch of amino acids vary and if they still serve a function remains unclear. The last 20 amino acids of the C-terminus of ureB are only represented in half of the compared sequences and accurate sequence conservation could not be determined in this part. This stretch contains a loop and a C-terminal helix **(Supp. Fig. 5)**. The ureC protein of the compared sequences has a shared sequence identity of 60.3%. All amino acids involved in catalysis are highly conserved **(Supp. Fig. 5)**. The ureA and ureB chains show lower conservation compared to ureC. They are not involved in catalysis but in scaffolding, so the differences could stem from their role in different types of oligomeric assembly **(Fig. 3)**.

**Figure 3.**
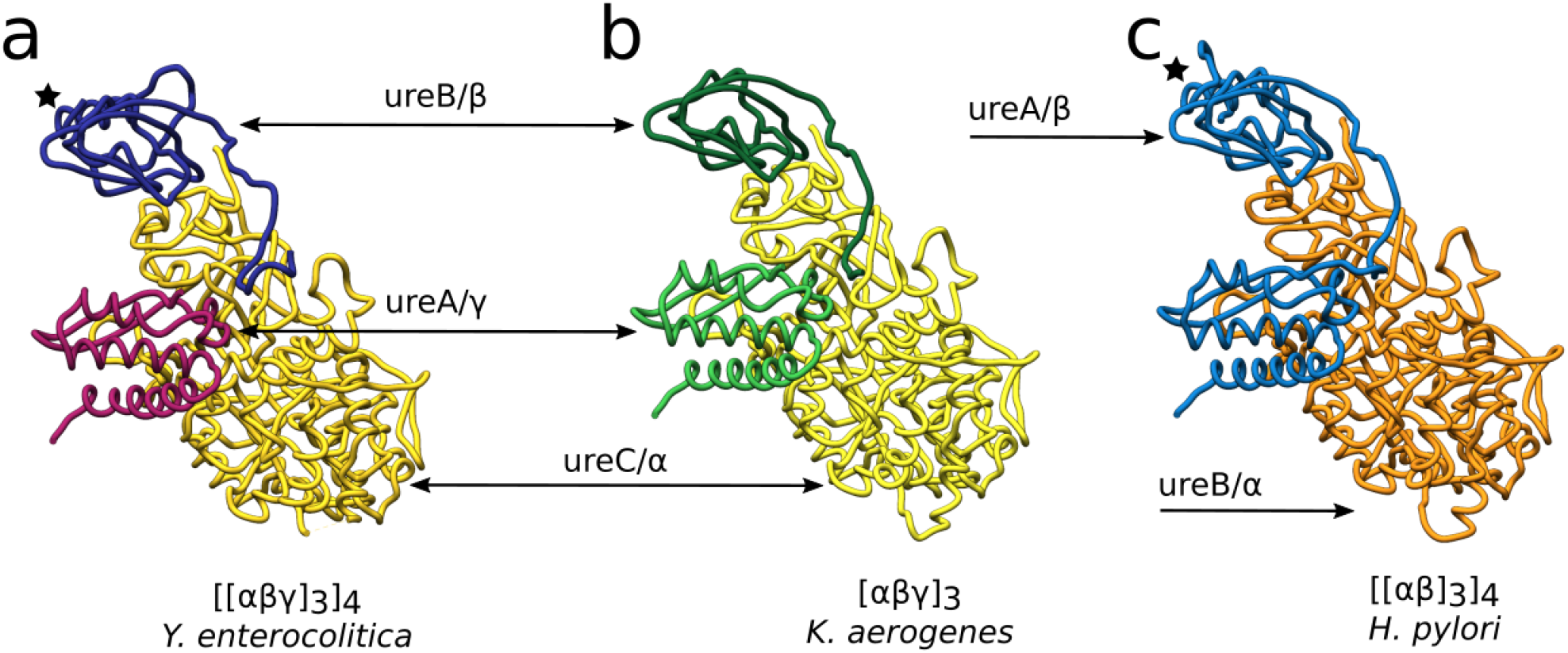
Comparison of *Y. enterocolitica* urease chain architecture with ureases with different modes of assembly from other pathogens. Hetero-trimers are shown in tube representation with each chain in the same colors of the sequences in **Fig. 1b**. The black star indicates the central helix of the β subunit. The *Y. enterocolitica* and *H. pylori* ureases form the same dodecameric assembly despite having different types of chain splitting, while *K. aerogenes* urease has the same type of chain splitting as in *Y. enterocolitica* but forms only a trimeric assembly.

To investigate this aspect further, we compared the presented structure to the ureases of *H. pylori, S. pasteurii* and *K. aerogenes*. Sequence identity scores among these ureases are provided in **Supp. Tab. 4**. The *H. pylori* urease is made up of two protein chains ureA (that contains the equivalent of ureA and ureB in *Y. enterocolitica)* and ureB (that is the equivalent of ureC in *Y. enterocolitica)* **(Fig. 1c and Fig. 3c)**. It assembles into a T-symmetric oligomer like in *Y. enterocolitica* and the crystal structure was solved to 3 Å. *S. pasteurii* and *K. aerogenes* ureases both assemble into a trimer from the hetero-trimeric unit **(Fig. 1c and Fig. 3b)**. *S. pasteurii* was solved in the presence of the inhibitor N-(n-Butyl)thiophosphoric Triamid (NBPT) to 1.28 Å (Mazzei *et al*., 2017). The crystal structure of *Klebsiella aerogenes* was solved to 1.9 Å as in absence of substrate or inhibitors (Jabri *et al*., 1995).

There are two main regions with high root mean square deviations (RMSDs) when comparing these three ureases to the *Y. enterocolitica* model **(Supp. Fig. 6 and Supp. Tab. 5)**. The first region with high deviation is the mobile flap, which opens and closes over the active site (residues 312 to 355 of ureC). The residues of its connecting loop could not be built with confidence in the cryo-EM model (residues 326 to 333 of ureC). The other region with large differences is on the edges of ureA and ureB where the interactions with the next protomer occur. The *H. pylori* assembly contains an additional C-terminal loop (residues 224-238 of ureA) after the top alpha helix (residues 206-223 of ureA). This helix (central helix) forms the three-fold axis of three neighboring trimers and the loop binds in a head-to-tail fashion to the next trimer forming the tetramer **(Fig. 3a,c)** (Ha *et al*., 2001). The core of the assembly is identical in its structure. For whole-chain superposition scores and RMSD values between the compared models please see **Supp. Tab. 5**.

In the dodecameric assembly seven different interfaces are formed between the hetero-trimers **(Fig. 4a,b and Supp. Fig. 7)**. Intra-trimer interactions occur between the three basic hetero-trimers in one assembled trimer, forming a three-fold symmetry axis **(Fig. 4a)**. The interactions between these trimers to form the tetramer then make up a different three-fold symmetry axis **(Fig. 4b)**. The three largest interfaces (interfaces 1-3) are formed intra-trimeric between ureC of one hetero-trimer and ureC, ureA and ureB of the next trimer **(Fig. 4a)**. The three ureA proteins make up the intra-trimer-core (first three-fold axis) with interface 4 **(Fig. 4c and Supp. Fig. 7)**. Comparison of the interface areas formed in the trimer assembly shows no substantial differences between the four organisms **(Supp. Fig. 7)**. Inter-trimer interfaces (interfaces 5, 6, 7) formed in the dodecameric *Y. enterocolitica* and *H. pylori* ureases have similar areas **(Fig. 4b and Supp. Fig. 7)**. Part of the interactions occur between ureB and ureC forming interfaces with each other (interface 4, 6). The other interaction is between the three ureB proteins and forms interface 7 and the inter-trimer-core (second three-fold axis) with their central helices **(Fig. 4b)**. *Y. enterocolitica* does not have the same oligomerization loop after the central helix proposed for *H. pylori*. However, there is a short loop before the central helix, which is extended in *Y. enterocolitica*. It binds into a pocket of ureC of the neighboring trimer in interface 6 **(Fig. 4e)**. These types of loops or extensions are missing from *S. pasteurii* and *K. aerogenes* ureB proteins. *S. pasteurii* has the central helix, but there is no extended loop before or after it **(Supp. Fig. 6b)**. *K. aerogenes* urease does not have a helix nor a loop in this region **(Fig. 3b)**. This suggests that the presence of oligomerization loops in ureB is crucial for determining the oligomeric state of the enzyme.

**Figure 4.**
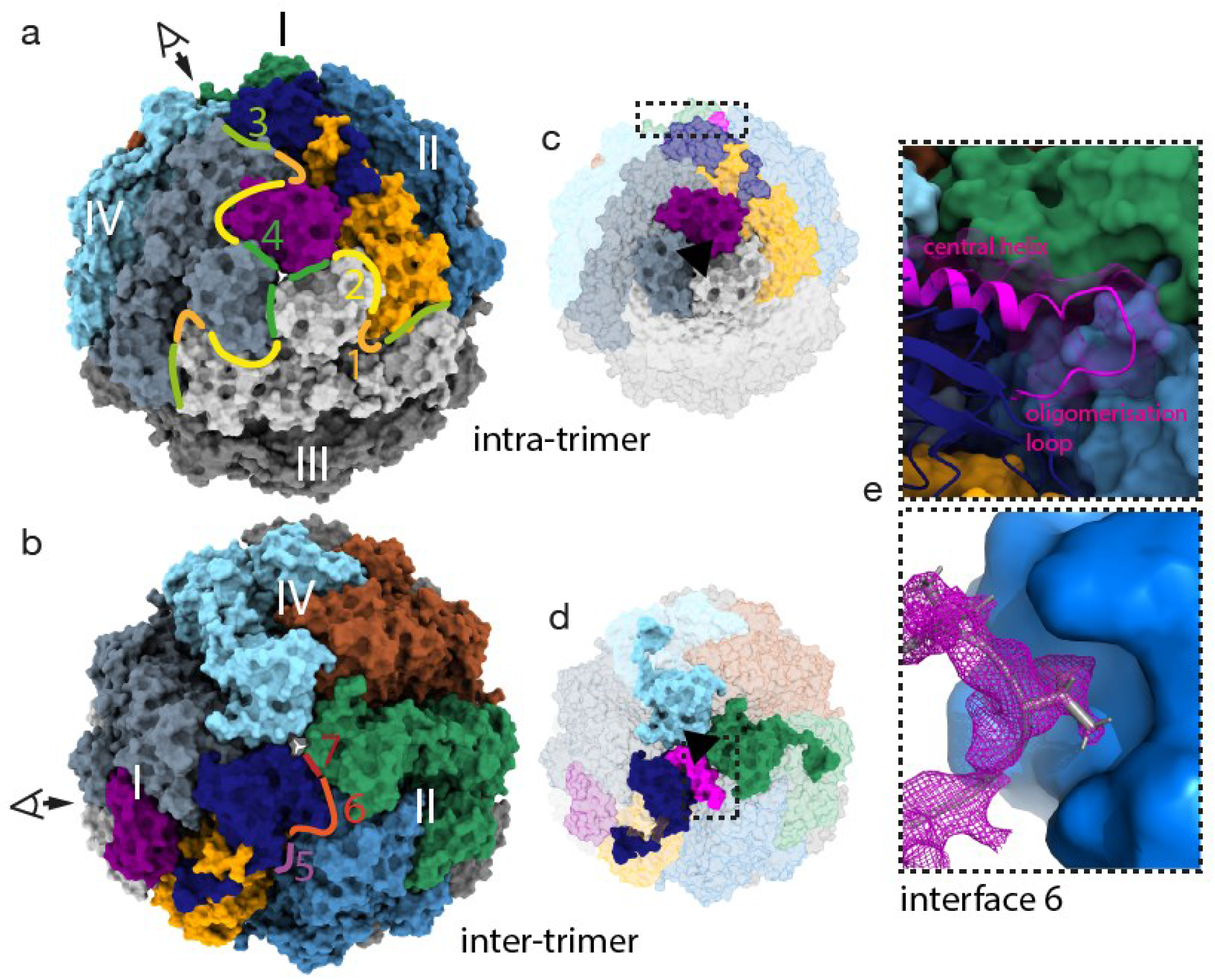
Interfaces in dodecameric assembly of *Y. enterocolitica*. **a)** surface model in front view of trimer of *Y. enterocolitica* urease with intra-trimeric interfaces 1-4 indicated with color-coded lines and numbers. **b)** Same model shown from the top (view indicated with eye) and the inter-trimeric interfaces. **c)** Front view with intra-trimeric-core highlighted and three-fold axis indicated with black triangle. Inset for e) in dashed box. **d)** same as c) but from the top view. **e)** Interface 6 with loop from ureB (magenta) binding into pocket of ureC of neighboring trimer. Upper inset shows ureB in cartoon and transparent surface and ureC in surface representation. Lower panel shows ureC as surface and ureB loop as cartoon with density.

The dodecameric holoenzyme structure of ureases might aid in stabilizing the protein at acidic pH, and in combination with 12 active sites producing ammonia enables the formation of a pH-neutralizing microenvironment around the assembly (Ha *et al*., 2001). This ensures the continued function of the enzyme and makes this type of oligomeric assembly essential to survival of *Y. enterocolitica* in the host. It is remarkable that *Y. enterocolitica* is the first organism outside the Helicobacteraceae family to have a known dodecameric urease. Considering the different subunit organization between these ureases, it raises the question of what particular events in the evolutionary history of *Y. enterocolitica* could have led to this type of assembly (Ligabue-Braun *et al*., 2013).

### The empty active site is filled with water

At the global resolution of 1.98 Å, detailed structural features can be observed. All throughout the highly resolved areas of the protein, salt bridges, backbone and side chain hydration and alternative side chain conformations can be visualized **(Supp. Fig. 8a-c)**. Furthermore, the high resolution allows for a detailed description of the nickel-metallo-center and the active site. The active site is located on the ureC protein at the edge of the hetero-trimer and is wedged in between the ureA and ureB proteins of the next hetero-trimer in the homo-trimeric assembly **(Fig. 5a)**.

**Figure 5.**
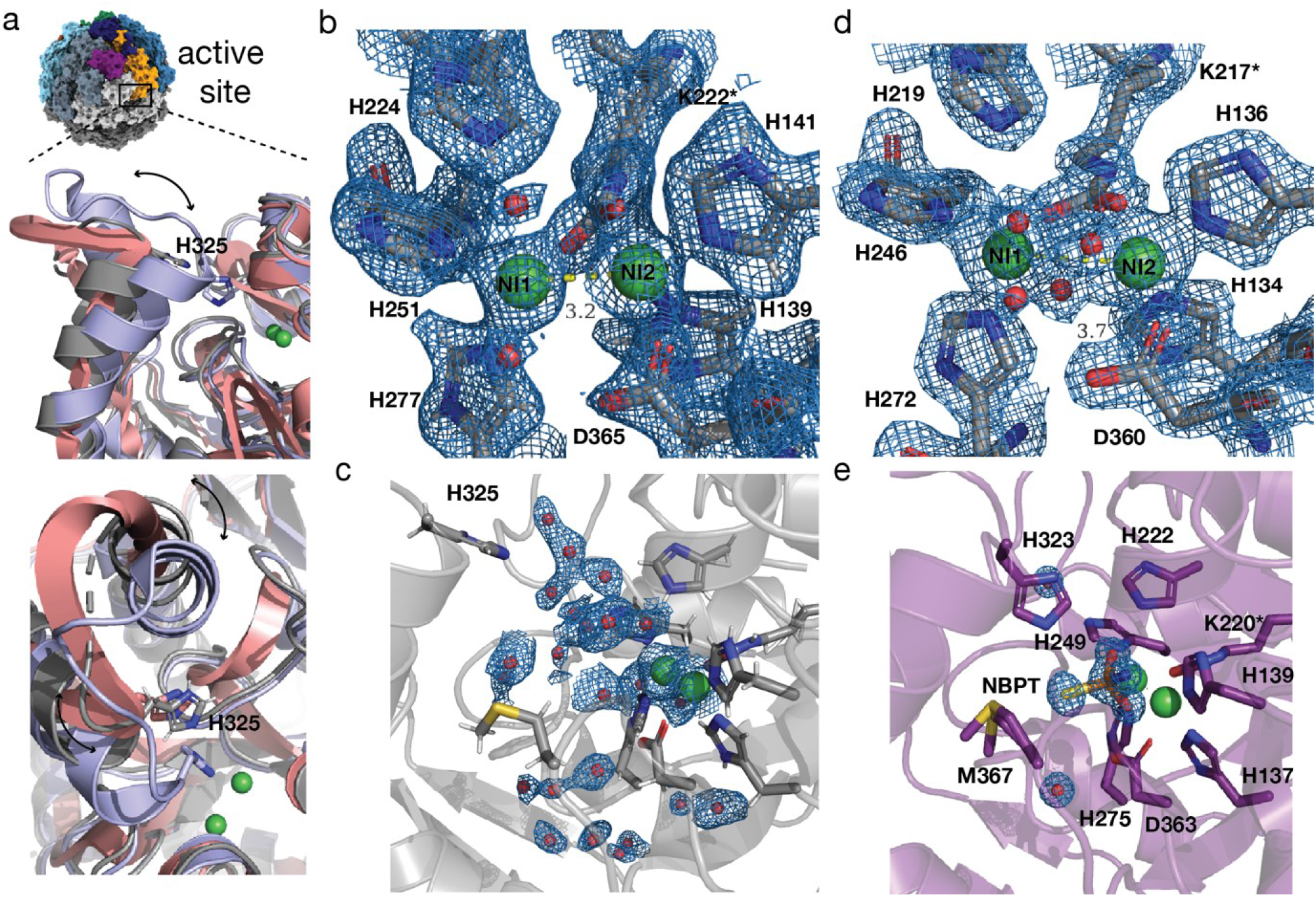
Active site of *Y. enterocolitica* urease. **a)** Overview of urease assembly with the active site location indicated. Inset shows in top panel side view of urease crystal structures from *K. aerogenes* mobile flap shown in open conformation in salmon (PDB: 2UBP) and in closed position as light purple (PDB: 3UBP). In gray the cryo-EM structure of *Y. enterocolitica* is overlaid and the green spheres represent the Ni^2+^ ions of the active site. Bottom panel shows top view of the three structures. Arrows indicate movement of helix and catalytic HIS325 is shown as stick. **b)** Model of active site residues and Ni^2+^ ions with the cryo-EM map of *Y. enterocolitica* at 1.98 Å nominal resolution. Yellow line indicates distance between Ni^2+^ ions in Å; **c)** shows the water molecules in the active site. **d)** Crystal structure of *K. aerogenes* urease at 1.9 Å resolution (PDB: 1EJW). Yellow line indicates distance between Ni^2+^ ions in Å. **e)** Crystal structure of *S. pasteurii* at 1.28 Å with inhibitor NBPT (PDB: 5OL4).

The catalysis of ammonia and carbamate from urea occurs in two steps **(Fig. 1a)**. Urea first interacts with the nickel ions through its carbonyl oxygen and amino nitrogens. The active site contains two Ni^2+^ ions which are coordinated by six different amino acids **(Fig. 5b)**. Both Ni^2+^ ions are coordinated by the carbamylated LYS222*. Ni(1) is additionally coordinated by HIS224, HIS251 and HIS277 and Ni(2) by HIS139, HIS141 and ASP365. Close to the active site is a methionine (MET369), which can be modelled in different alternative conformations. One conformation could potentially reach the active site. There is no described function for this amino acid **(Fig. 5c and Supp. Fig. 9)**.

The active site is protected by a helix-turn-helix motif, called the mobile-flap. Its function is to coordinate the access of substrate to the catalytic site and the release of the product from it (Maroney & Ciurli, 2014). After closing of the mobile flap, a proton is transferred from a water molecule to the other amine in a nucleophilic attack on the carbonyl carbon of urea. A conserved histidine on the mobile flap (HIS325) is essential for catalysis by possibly acting as a general acid and aiding in deprotonation **(Fig. 5a)** (Maroney & Ciurli, 2014; Kappaun *et al*., 2018). By closing of the mobile flap the HIS325 moves closer to the active site, also stabilizing urea in the active site pocket (Maroney & Ciurli, 2014). Flap opening then releases ammonia and carbamate, where the latter spontaneously hydrolyses into another molecule of ammonia and bicarbonate. The mobile flap of the cryo-EM structure presented here is in an open position, which is explained by the absence of substrate or inhibitors in the sample **(Fig. 5a)**. Coordinated water molecules can be seen in the empty pocket of the active site, which do not only form hydrogen bonds with side chains or the protein backbone, but also with each other constituting a hydration network **(Supp. Fig. 8d)**.

The resolution in the active site is sufficient for complete atomic description of the coordinated Ni^2+^ ions, including the carbamylated lysine. The protonation states of the active site residues are also represented in the map **(Fig. 5b)**. One of the hydroxide molecules in the active site is essential as it performs the nucleophilic attack on urea while other molecules are displaced by urea and the closing of the mobile flap (Kappaun *et al*., 2018). Comparison to the crystal structure of *K. aerogenes* of similar nominal resolution (1.9 Å) shows differences in the visualization of these features. This crystal structure was solved in absence of inhibitors or substrate such that the active site is also empty and the mobile flap in an open conformation **(Fig. 5a)**. The details of the map provides finer details around the Ni^2+^ ions in the cryo-EM map than the crystallographic data. The protonation of the histidines is clearly visible in the cryo-EM density **(Fig. 5b)**. The positions of the side chains and the Ni^2+^ ions in the active site are very similar to the *Y. enterocolitica* urease structure with a RMSD of 0.270 Å **(Supp. Tab. 5)**. The *S. pasteurii* crystal structure was solved in presence of the inhibitor NBPT which displaces the essential water molecules needed for the reaction from the active site. The closing of the mobile flap displaces the rest of the waters and brings the catalytic HIS323 closer to the active site. The tight packing of side chains prevents urea from entering the active site, efficiently blocking it **(Fig. 5b,e)**. The active site residues and Ni^2+^ ions have a RMSD of 0.293 Å between *S. pasteurii* and *Y. enterocolitica*.

### Nickel atoms come closer together

The distance between the Ni^2+^ ions is 3.7 Å in X-ray structures of *K. aerogenes* and *S. pasteurii*, but only 3.2 Å in the *Y. enterocolitica* cryo-EM model **(Fig. 5b,d)**. Short distances of 3.1-3.3 Å were described for *S. pasteurii* and *K. aerogenes* at high resolutions for structures in presence of β-Mercaptoethanol (β-ME) (Benini *et al*., 1998). Knowing that metallic cores are particularly sensitive to radiation (Yano *et al*., 2005), we tried to determine the extent to which radiation damage can explain the shorter distance between the Ni^2+^ ions. For this purpose, we generated per-frame reconstructions for the first 25 frames of our data collection and refined the model on each of them (see **Methods**) and measured the distances between the residues involved in ion coordination, shown in **Fig. 6**. At the beginning of the exposure, in which the frames contribute more to the full reconstruction due to dose-weighting (Grant & Grigorieff, 2015; Zivanov *et al*., 2019a), there is a trend of the ions coming closer together (**Fig. 6a-i**). While we cannot determine exactly how this arises from radiation damage, it is likely a result of several interactions in the active site changing simultaneously along the exposure. For example, both Ni(1) and Ni(2) tend to come closer to the carbamylated LYS222 (**Fig. 6a-iv,v**) as ASP365 vanishes (**Fig. 6a-vi**), which can be seen in the **Supp. Mov. 2**. Aspartic acid is known to have its side chain damaged very early on (Hattne *et al*., 2018). The dynamic interplay between residues along the exposure (**Fig. 6b**) is likely due to the different rates at which specific types of bonds and residues are damaged (Fromm *et al*., 2015): first negatively charged residues, then positively charged ones followed by aromatic side chains, as also observed in **Supp. Mov. 2**. However, the later part of the exposure must be interpreted with caution, as atomic coordinates become less reliable, which is verified by the overall increase in B-factors in **Fig. 6c** and the error bars in **Fig. 6a**.

**Figure 6.**
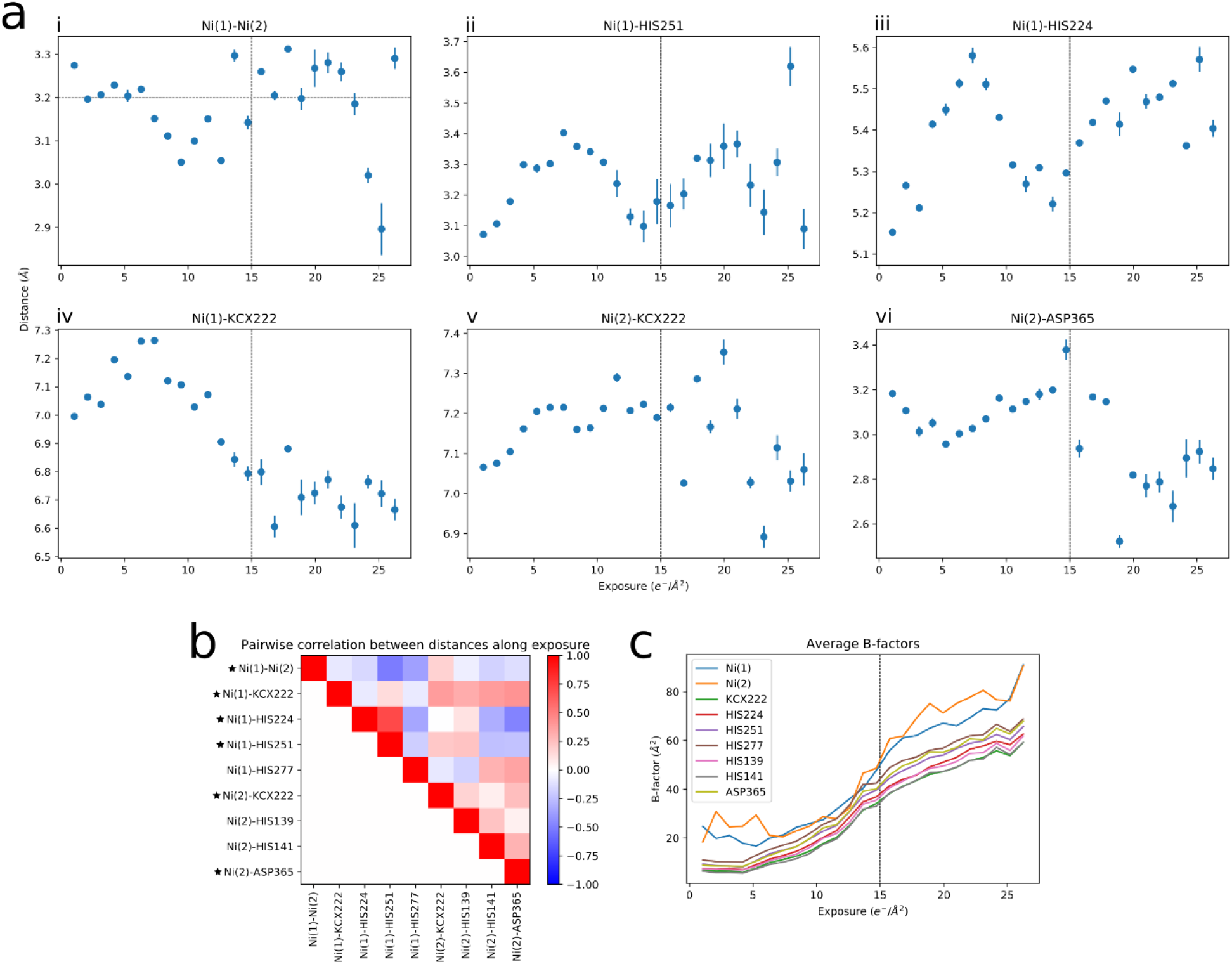
Radiation damage affects the distance between residues in the active site. **a)** Distances between the Ni^2+^ ions and selected residues involved in their coordination are plotted against the accumulated exposure. For each reconstruction calculated along the exposure, the model was refined, and distances measured. Dots indicate the average and error bars show +/- one standard deviation across five refinement runs with different random seeds. Horizontal dashed line in panel a-i) shows the distance in the model obtained from the full reconstruction with all frames. Vertical dashed lines show approximately the exposure at which the density for charged residues completely vanishes (see **Supp. Mov. 2**). **b)** Correlation coefficients between distance changes along the exposure for selected residues involved in ion coordination. Distance plots shown in a) are indicated with a star. **c)** Average B-factors of selected residues plotted against the accumulated exposure.

## Conclusion

Large urease assemblies have been historically difficult to study by X-ray crystallography (Ha *et al*., 2001). We have determined the structure of a dodecameric urease assembly, a metalloenzyme from the pathogen *Y. enterocolitica* at an overall resolution of 1.98 Å using cryo-EM. The collection of datasets with and without beam-image shift demonstrates the advantages of using this feature of modern TEMs and invites further investigations on the behavior of optical aberrations and specimen drift.

Our results demonstrate the feasibility of cryo-EM as a technique for obtaining structures of clinically relevant enzymes with sufficient quality for *de novo* model building and drug design. The cryo-EM map has allowed a detailed description of the active site and the oligomeric assembly. More specifically, we could observe the putative oligomerization loop that enables the dodecameric assembly, which was hypothesized to be responsible for the enhanced survival of *Y. enterocolitica* in highly acidic environments (Young *et al*., 1996). This urease is the first outside the Helicobacteraceae family, and therefore without an αβ subunit organization, to have a dodecameric assembly reported. What evolutionary events have led to this intriguing combination of subunit organization and quaternary structure are unknown.

Furthermore, in comparison to the *K. aerogenes* structure, which is at approximately the same nominal resolution, the cryo-EM map offers an improved representation of protons and Ni^2+^ ions. A possible explanation is that the error in the phases derived in the X-ray structure determination grows faster towards the limit of observed diffraction. Another aspect to be considered is that X-rays and electrons probe different properties of matter, respectively the electron density and the integrated Coulomb potential. Our results prompt a more detailed investigation of these effects and how they affect the representation of features at high resolution.

Finally, we noticed that radiation damage can partially explain the shorter distance observed between the nickel atoms in the active site. Given that ions and charged residues are damaged very early on in the exposure (Yano *et al*., 2005; Hattne *et al*., 2018), this effect cannot be neglected in structures derived from cryo-EM reconstructions. Novel direct electron detectors with higher frame rates may allow time-resolved experiments to investigate these effects in more detail.

## Supporting information

Supplementary Movie 1. Overview and feature highlights of the Yersinia enterocolitica urease cryo-EM structure.

Supplementary Movie 2. Morphing between maps and models obtained from different sets of frames along the exposure.

## Data availability

The model has been deposited at the PDB under accession code 6YL3. The map has been deposited at the EMDB under accession code EMD-10835. Raw electron microscopy data is deposited in EMPIAR, accession code EMPIAR-10389.

## Acknowledgments

The authors would like to thank L. Kovacik and K. Goldie for assistance in data collection and S. Klumpe, A. Nunes-Alves, R. Ligabue-Braun and T. Nakane for discussions. Cryo-EM data processing calculations were performed at sciCORE (http://scicore.unibas.ch/) scientific computing center at the University of Basel. R.D.R. and L.A. acknowledge funding from the Fellowships for Excellence program sponsored by the Werner-Siemens Foundation and the University of Basel. This work was in part supported by the Swiss National Science Foundation (grants 177195 and 185544, NCCR TransCure).

## Author Contributions

R.D.R. and R.A. performed the cryo-EM experiments and data analysis. L.A. built and analyzed the atomic model. R.P.J. performed X-ray crystallography experiments. M.A.M. expressed and purified the protein. P.R. prepared and screened EM samples. J.Z. performed the higher-order aberration corrections and analysis. T.S., T.M. and H.S. initiated and supervised the project. R.D.R, L.A., R.A., T.M. and H.S. wrote the manuscript with assistance from all authors.

## Methods

### Protein expression and purification

The *Y. enterocolitica* urease was purified for cryo-electron microscopy according to the protocol of (Rokita *et al*., 2000). The strain was precultured overnight at 37°C for 18 hours in a medium containing 37 g/l of brain/heart infusion (Oxoid, CM0225), 50 μg/ml streptomycin sulfate (Applichem, A1852.0100), 35 μg/ml nalidixic acid (Applichem, A1894.0025), 50 μg/ml meso-diaminopimelic acid (Sigma, D1377) and 100 μM nickel(II) chloride hexahydrate (Sigma, N6136). 6 x 600ml of expression cultures were inoculated at OD of 1 at 28°C for 23 hours. Cells were harvested by centrifugation and the cell pellet resuspended in 0.15M NaCl, 50mM Tris pH 8.0. The cell lysate was applied directly to a Sephacryl S-300 HR 26/60 column equilibrated with 150 mM NaCl, 50mM TrisHCl pH 8.0. The active fractions as identified by a phenol-hypochlorite assay (Weatherburn, 1967) were buffer-exchanged to 50mM Tris pH 7.0 within a centrifugal filter unit (Sartorius, Vivaspin MWCO 50kDa) and applied to a Mono Q HR 5/5 column pre-equilibrated with 50mM TrisHCl, pH 7.0. The protein was eluted in 50mM Tris pH 7.0 by a gradient to 1M NaCl, concentrated on a centrifugal filter unit (Sartorius, Vivaspin MWCO 50 kDa) and purified by SEC as before. The purity of the urease sample of the two preparations was verified on a 4%/12% SDS-PAGE and by mass spectroscopy.

### Sample preparation

Approximately 3 μl of the 0.39 mg/ml urease solution were applied to glow-discharged Quantifoil holey carbon grids. After 3-second blotting, the grids were flash-frozen in liquid ethane, using a FEI Vitrobot IV (Thermo Fisher Scientific) with the environmental chamber set at 90% humidity and 20 °C temperature.

### Data acquisition

Cryo-EM data were collected on a FEI Titan Krios (Thermo Fisher Scientific) transmission electron microscope, operated at 300 kV and equipped with a Quantum-LS imaging energy filter (GIF, 20 eV zero loss energy window; Gatan Inc.) and a K2 Summit direct electron detector (Gatan Inc.) operated in dose fractionation mode. Data acquisition was controlled by the SerialEM (Schorb *et al*., 2019) software, performed in counting mode, with a 42 e^-^/Å^2^ total exposure fractioned into 40 frames over 8 seconds. The physical pixel size was 0.639 Å at the sample level. The data was pre-processed via the FOCUS package (Biyani *et al*., 2017), including drift-correction and dose-weighting using MotionCor2 (Zheng *et al*., 2017) (grouping every 5 frames and using 3×3 tiles) and CTF estimation using CTFFIND4 (Rohou & Grigorieff, 2015) (using information between 30 Å and 5 Å from the movie stacks). With these settings, we collected two datasets: one using beam-image shift (Cheng *et al*., 2018), with three shots per grid hole, comprising 2,243 movies, and a second one taking a single shot per hole, with 2,252 movies. A summary of data collection information is given in the **Supp. Tab. 1**.

### Image processing

The two datasets were initially processed separately as shown in the flowchart of **Supp. Fig. 2**. We excluded all movies whose resolution of CTF fitting was worse than 4 Å according to CTFFIND4, leaving 2,197 movies in the multi-shot dataset or 2,115 in the single-shot dataset for further processing. Using the template-free LoG-picker algorithm (Zivanov *et al*., 2018) we picked an initial set of 157,699 particle coordinates on the multi-shot dataset. These particles were extracted and subjected to one round of 2D classification with the aim of removing “bad” or false-positive particles. Best results in 2D classification were observed when enabling the RELION option “Ignore CTFs until first peak?”. Selecting only the classes displaying views of urease with high resolution features, a new subset containing 60,271 particles was obtained. Using this subset, a first 3D map was obtained by the *ab initio* stochastic gradient descent (SGD) algorithm (Zivanov *et al*., 2018; Punjani *et al*., 2017) with and without tetrahedral symmetry imposed. The symmetric map was consistent with previously determined structures of ureases (Arnold *et al*., 2016; Ha *et al*., 2001). The particles were then subjected to 3D refinement using the map from the *ab initio* procedure as starting reference, resulting in a map at 2.6 Å resolution. We then generated new templates for particle picking by low-pass filtering the unsharpened map from this first 3D refinement to 20 Å and calculating evenly oriented 2D projections from it. These templates were then used for picking with Gautomatch (K. Zhang, http://www.mrc-lmb.cam.ac.uk/kzhang/), detecting 107,399 particle coordinates on the multi-shot dataset or 87,204 particle coordinates on the single-shot dataset. Visual inspection of randomly selected micrographs indicated this set of coordinates was better than that previously found by the LoG-picker, in the sense that it contained fewer false positives and more true particles. The newly extracted particles were then subjected to 2D and 3D classification procedures to get rid of false positive, damaged or broken particles, which yielded cleaner subsets with 62,884 (multi-shot) or 51,173 (single-shot) particles. Using the current best map from the multi-shot dataset as a starting reference, we then performed masked 3D refinements on the two datasets separately, interleaved with rounds of CTF refinement and Bayesian particle polishing (Zivanov *et al*., 2019a). More specifically, we refined defocus *per particle*, astigmatism *per micrograph* and beam tilt *globally* in CTF refinement. In the multi-shot dataset, each of the three relative “shooting targets” were assigned a different class for separate beam tilt refinement. The parameters for Bayesian particle polishing were trained separately on ∼5,000 particles from each dataset at this stage. Each dataset yielded refined maps at 2.10 Å (multi-shot) and 2.20 Å (single-shot) resolution. Best results, however, were obtained when merging the particles picked by template-matching on each dataset (194,603 particles in total) and processing them altogether. After 2D classification, 141,069 particles remained **(Supp. Fig. 1b),** and after 3D classification, there were 119,020 particles **(Supp. Fig. 1c)**. CTF refinement was then performed using four beam tilt classes, with the particles from the single-shot dataset belonging to a new, fourth class **(Supp. Tab. 2)**. Defocus and astigmatism were both refined *per particle* this time, resulting in a map resolution of 2.05 Å. We compared polishing the full merged dataset at once and each beam tilt class separately, to verify if there were differences in the patterns of particle motion. For training the polishing parameters ∼10,000 particles were used in each case this time. Although we did observe different statistics of particle motion **(Supp. Tab. 3)**, resolution and overall quality of the map did not improve further by performing either merged or separate polishing of the different shots. Finally, correction of third-order aberrations in RELION-3.1 (Zivanov *et al*., 2019b) **(Supp. Fig. 3)** followed by local 3D refinement resulted in a global map resolution of 1.98 Å. The angular distribution efficiency was estimated using the program cryoEF (Naydenova & Russo, 2017).

All resolution estimates reported were obtained by considering the 0.143 threshold (Rosenthal & Henderson, 2003) on the Fourier shell correlation (FSC) curve (Harauz & van Heel, 1986) between independently refined half-maps (Scheres & Chen, 2012). A solvent-excluding mask was generated by low-pass filtering the maps to 12 Å, binarizing the filtered map and adding a soft edge consisting of a cosine-shaped falloff to zero. The FSC curve was corrected for artificial correlations introduced by the mask (Chen *et al*., 2013). Local resolution was estimated using the approach implemented in RELION (Cardone *et al*., 2013).

### Model building, refinement and analysis

After refinement of the map to high resolution it had to be flipped in UCSF Chimera (Pettersen *et al*., 2004) to match the correct handedness. The biological assembly from the crystal structure of *Y. enterocolitica* urease **(Supp. Note 1)** was rigid-body fitted to the map in Chimera. The non-crystallographic symmetry (NCS) was calculated with PHENIX v1.17 (Adams *et al*., 2002) from the crystal structure. Only using the hetero-trimer of the three proteins, backbone and side chains were built, corrected or confirmed in Coot (Emsley & Cowtan, 2004). After several rounds of manual refinement of the model in Coot, applying NCS and real-space refinement in PHENIX, the model comprised side chains of residues 1-100 of ureA, 31-162 of ureB and 2-327, 335-572 of ureC. Residues 328-334 are disordered and could not be modeled with confidence. NCS constraints were not used during final refinements as to include alternative side chain conformations. Waters were built manually and refined in PHENIX and were added to the closest chain by the program phenix.sort_hetatoms. The quality of the refinement was assessed by cryo-EM Validation tool **(Table 1)** (Afonine, Klaholz *et al*., 2018).

### Structure analysis

The electrostatic potential was calculated using the APBS plugin in PyMOL (Jurrus et al, 2018). The ConSurf server (Berezin *et al*., 2004; Ashkenazy *et al*., 2016) was used to find 150 sequences per urease protein for alignment with ClustalW (Madeira *et al*., 2019) and calculate conservation per residue. The “sample the list of homologs” option was used to get a diverse representation across all species. The sequence identity of the 150 sequences was determined using BLSM62 in Geneious (Kearse *et al*., 2012). The PDBePISA server (https://www.ebi.ac.uk/pdbe/pisa/) was used to find and calculate interface areas (Krissinel & Henrick, 2007). Figures were created using UCSF Chimera (Pettersen *et al*., 2004), ChimeraX (Goddard *et al*., 2018), PyMol (Schrödinger), Inkscape, Adobe Illustrator and Adobe Photoshop.

### Radiation damage analysis

Bayesian particle polishing in RELION (Zivanov *et al*., 2019a, 2018) was carried out on a sliding-window basis along the exposure, including 5 frames at a time, starting from frame 1 up to frame 25. Half-set reconstructions were then created from each polished particle stack and post-processed using the same mask as that applied to the reconstruction from all frames. On each post-processed reconstruction, real space refinement of chain C (α-subunit containing the active site) from the full reconstruction was carried out in PHENIX (Afonine, Poon *et al*., 2018) for 5 macro-cycles. This procedure was repeated 5 times with different random seeds. Distance between residues in the resulting refined models were calculated using BioPython (Cock *et al*., 2009) and plotted using the NumPy (https://www.numpy.org) and Matplotlib (https://www.matplotlib.org) Python modules.

## Supplementary Information

### Supplementary Figures

**Supplementary Figure 1.**
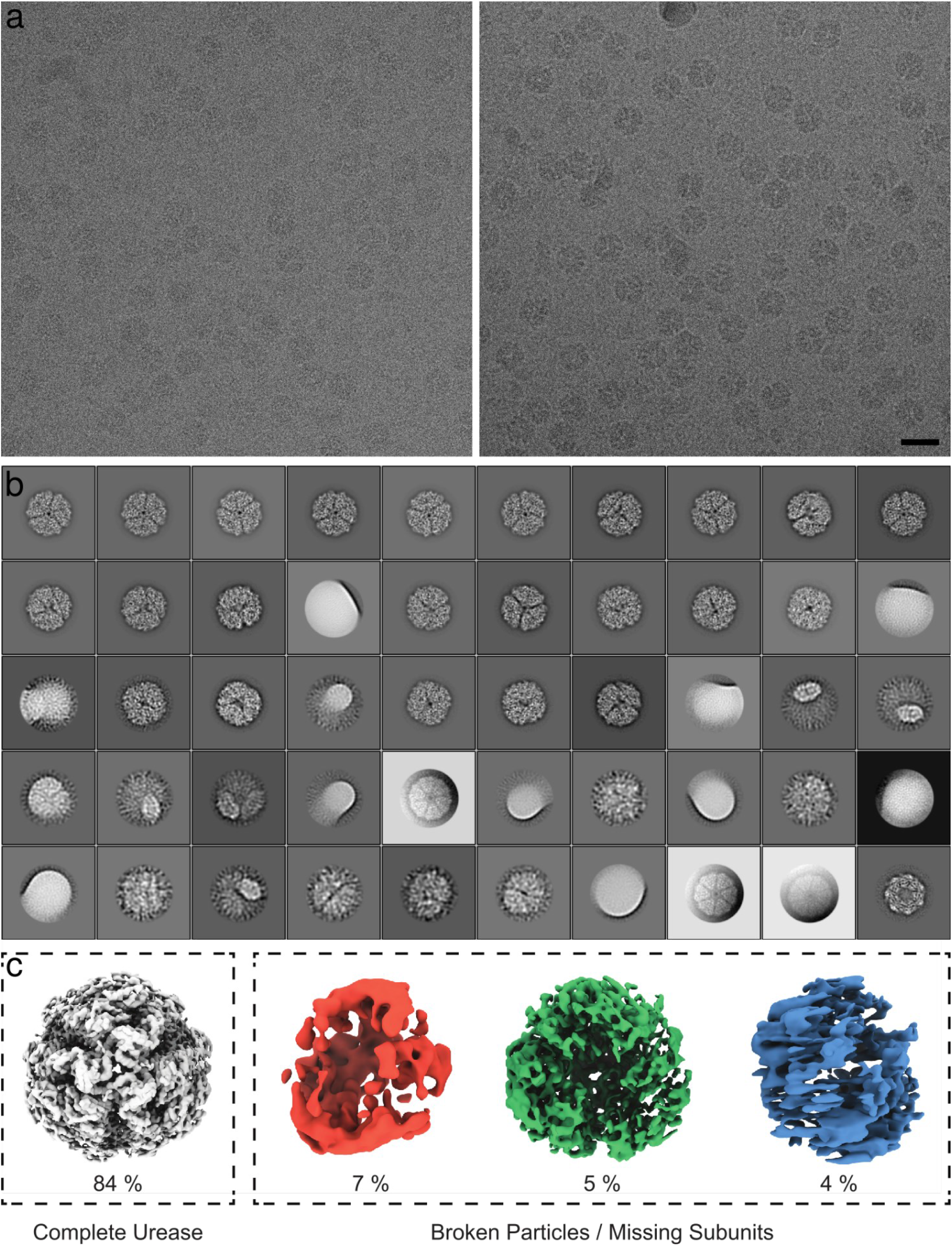
Cryo-EM data of *Y. enterocolitica* urease. **a)** Two representative micrographs from the dataset, acquired at −0.64 and −1.36 μm defocus, respectively. Scale bar: 200 Å. **b)** 2D class averages obtained from 194’603 particles in the merged dataset. These averages were obtained with RELION’s “Ignore CTF until first peak” option enabled and are sorted by decreasing order of number of particles in each class. Views of urease with missing subunits are observed. The bottom right average shows a contamination by GroEL. **c)** 3D class averages obtained from 141’069 particles in the merged dataset. These averages were obtained without symmetry imposition in RELION. The 3D classes are sorted by the indicated fraction of particles assigned to it. The first class is a complete dodecameric urease assembly, while the other classes represent urease structures with at least one trimer missing from the tetrahedron.

**Supplementary Figure 2.**
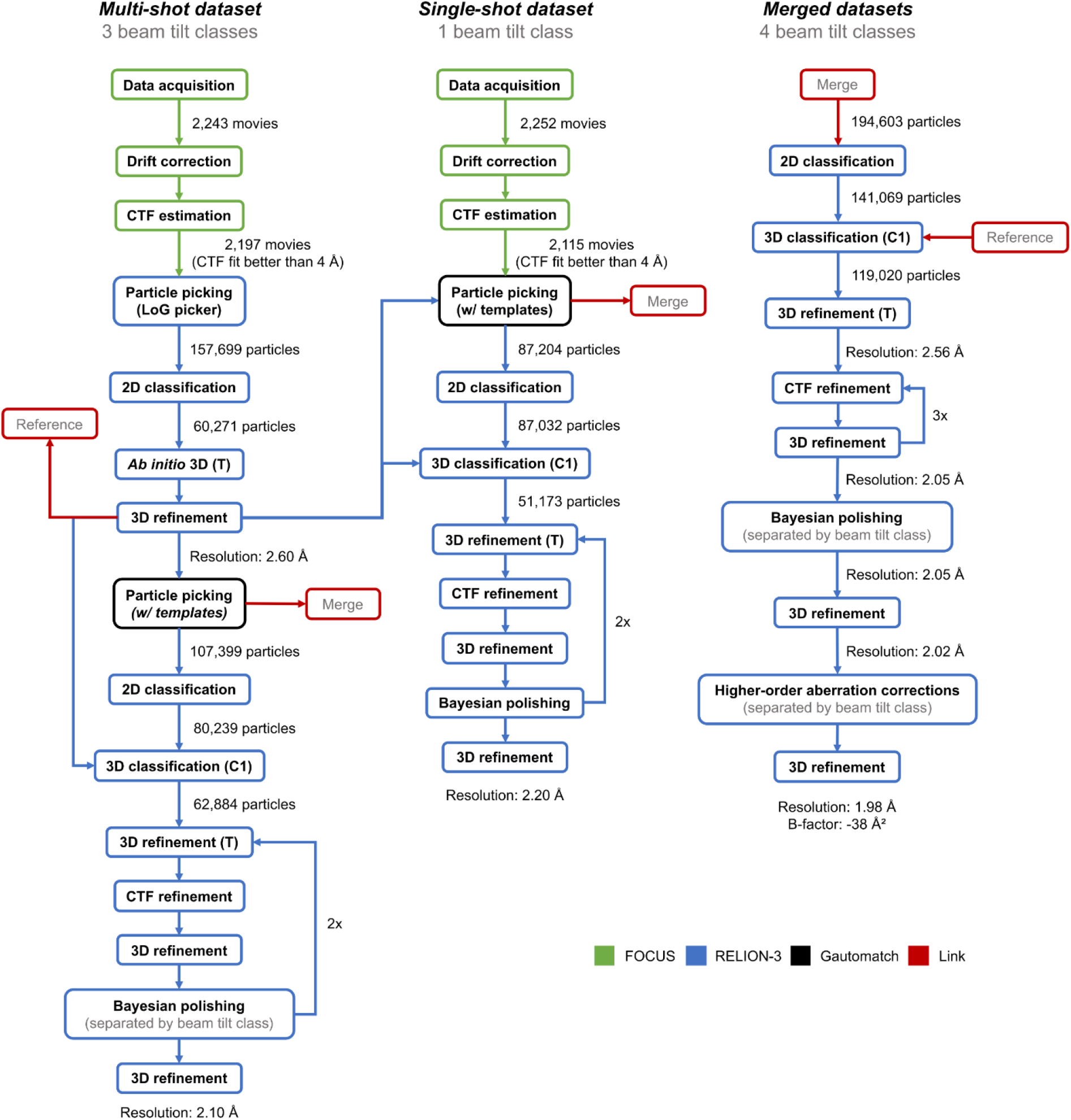
Data processing flowchart for the urease cryo-EM map. Masking and postprocessing jobs have been omitted for clarity. All CTF refinement jobs included beam tilt and per-particle defocus refinement (see **Methods** for details). All resolution estimates given correspond to the corrected FSC curves between masked half-maps obtained from postprocessing jobs in RELION.

**Supplementary Figure 3.**
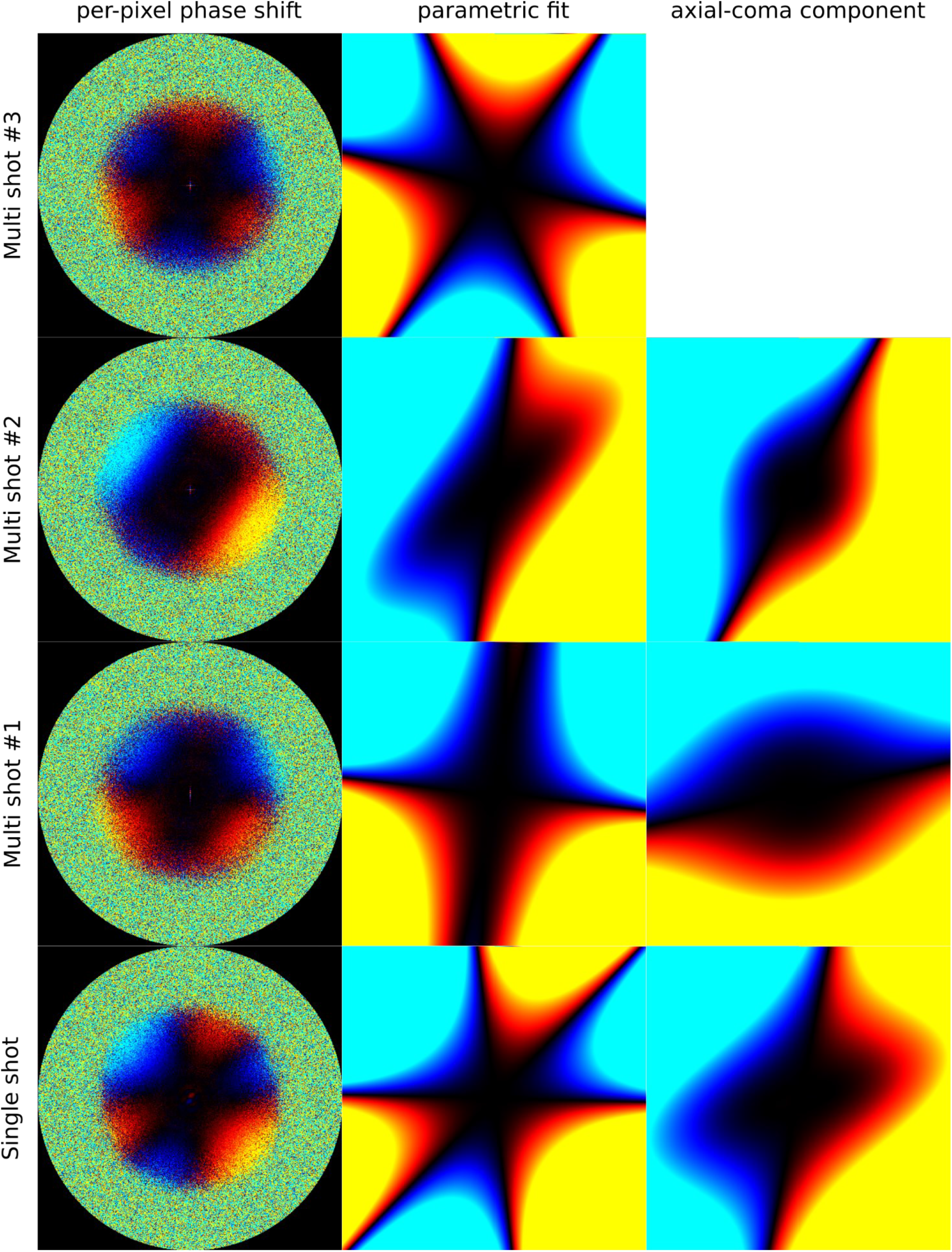
Fits of the anti-symmetrical aberrations arising at the four different beam-shift positions. The left column shows phase shifts measured independently for each Fourier pixel, while the center column shows their parametric fits using third-order Zernike polynomials. The first position (top row, the third beam-shifted multi-shot) corresponds to an essentially untilted beam **(Supp. Tab. 2)**, while the other two multishots and the single-shot dataset exhibit tilts to different extents. Note that even the untilted dataset shows a significant trefoil aberration. In the right column, the parametric fit of the first position has been subtracted, yielding residuals roughly consistent with the axial coma produced by a tilted beam.

**Supplementary Figure 4.**
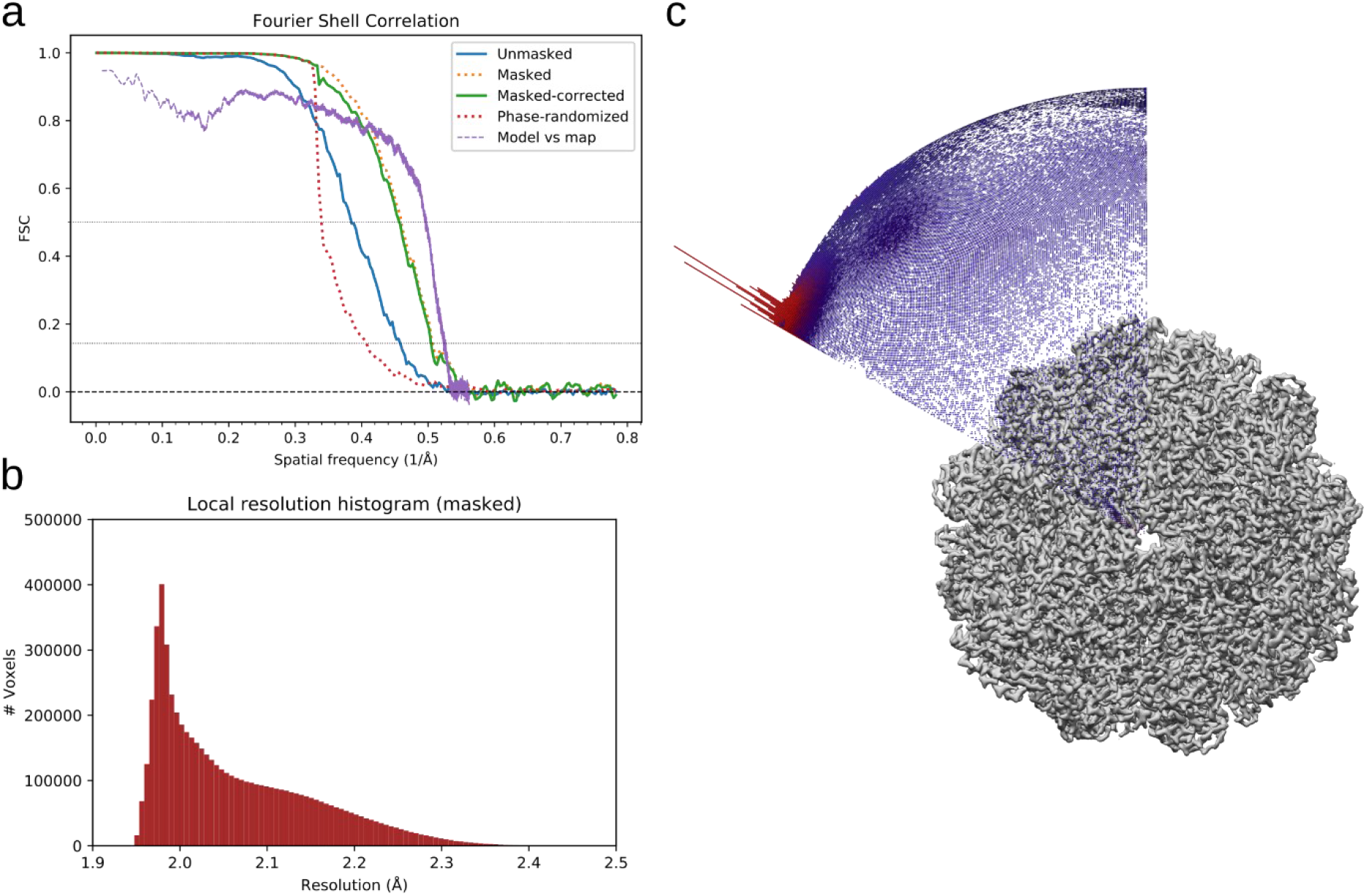
Resolution estimates of the urease cryo-EM map. **a)** FSC curves between the half-maps when unmasked (solid blue line), masked (dashed orange), corrected by high-resolution noise substitution after masking (solid green), phase-randomized (dashed red) and between the atomic model and the full experimental map (dashed violet). **b)** Histogram of local resolution assigned to each voxel. **c)** Angular distribution of particles in the urease cryo-EM reconstruction overlaid on the unsharpened map.

**Supplementary Figure 5.**
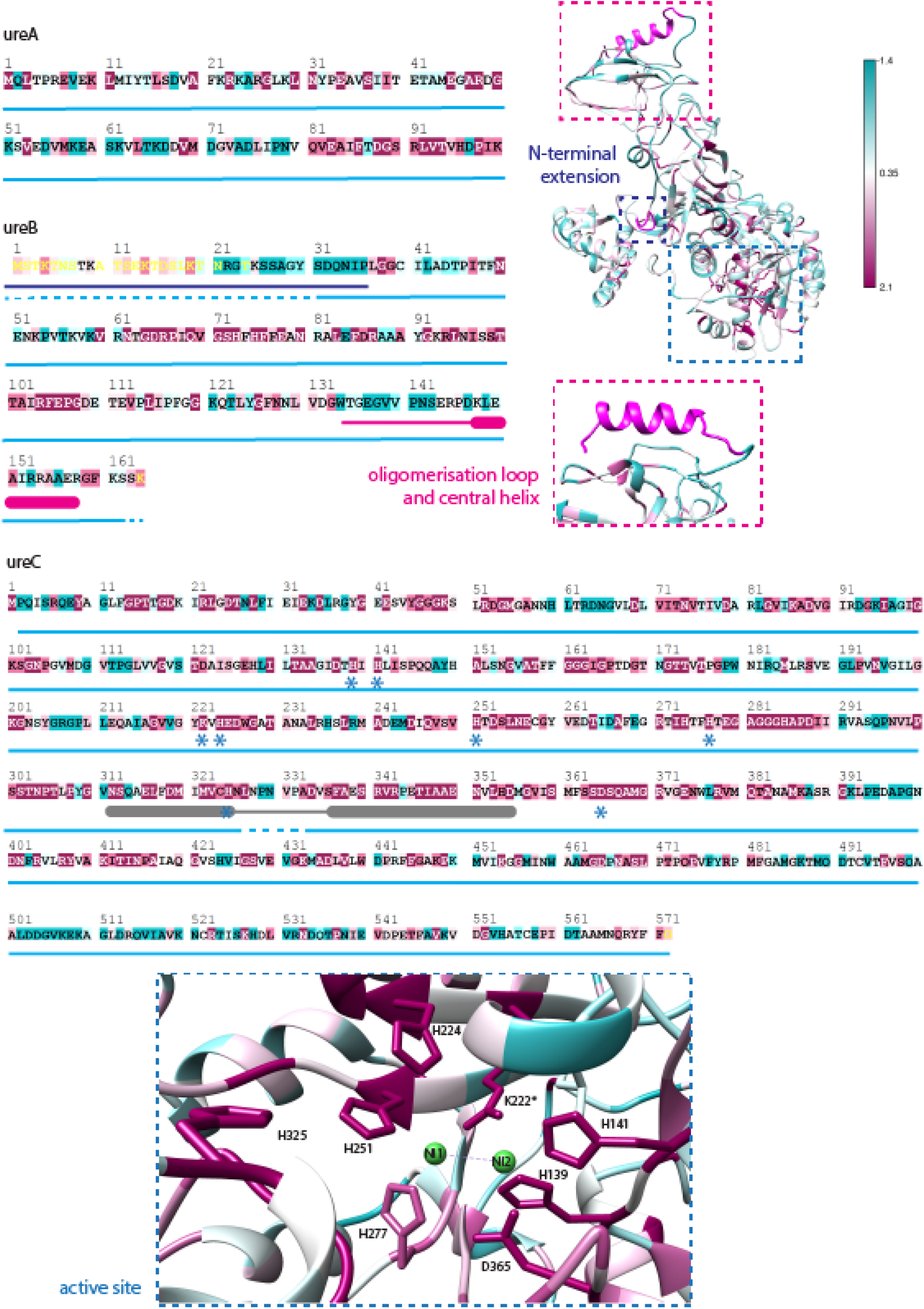
Alignment of 150 ureA, ureB and ureC sequences, chosen by sampling from all homologs found for each protein. ClustalW was used for sequence alignment and the ConSurf server for conservation analysis. Conservation is shown on a gradient from dark purple to white to turquoise (arbitrary units). The light blue bar indicates model completeness. Regions of interest are highlighted: The N-terminal extension in dark blue, the oligomerization loop and its following helix are indicated in magenta, the mobile flap is indicated in grey and the blue asterix indicates residues belonging to the active site.

**Supplementary Figure 6.**
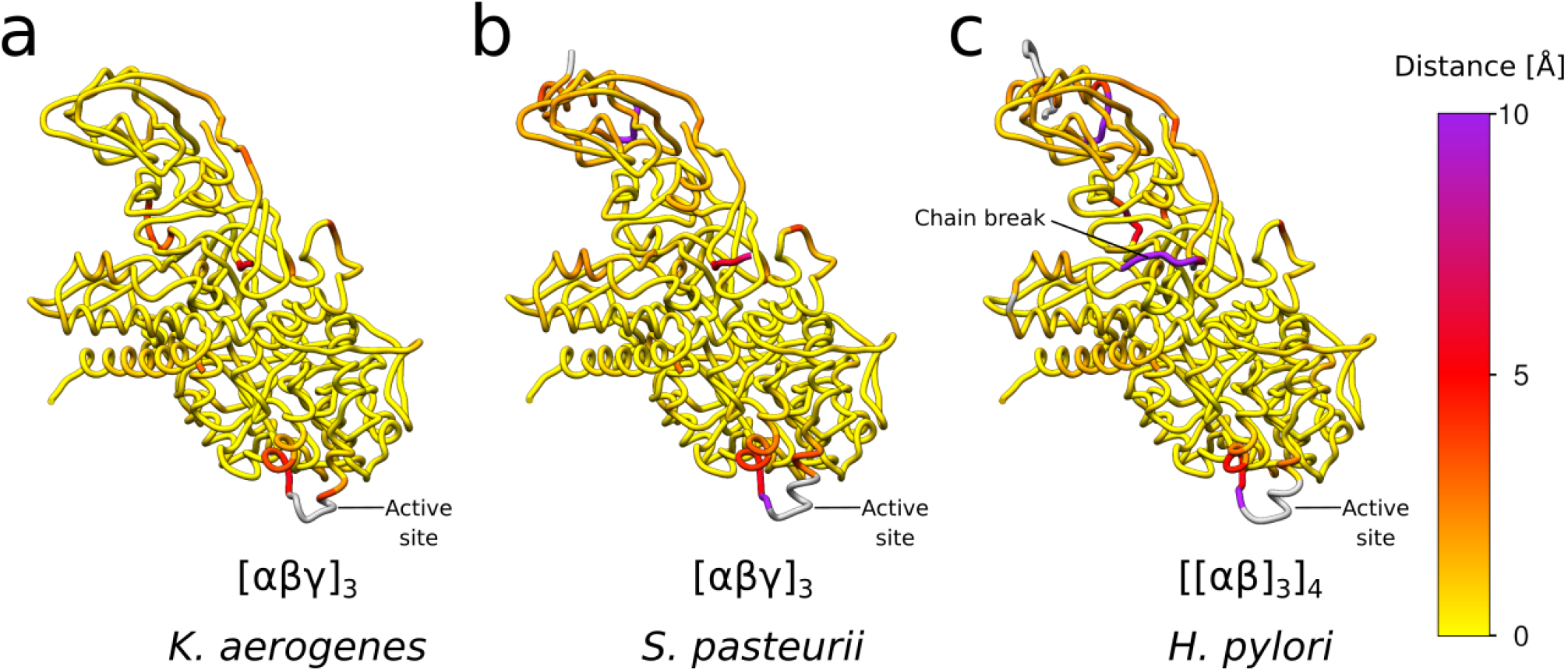
Distance comparison between residues of *Y. enterocolitica* urease against ureases with different modes of assembly. Tubes are colored by the pairwise distance of Cα atoms to the corresponding residue in *Y. enterocolitica* urease. Residues without equivalence after sequence alignment are shown in gray color. Segments with particularly high deviations are indicated.

**Supplementary Figure 7.**
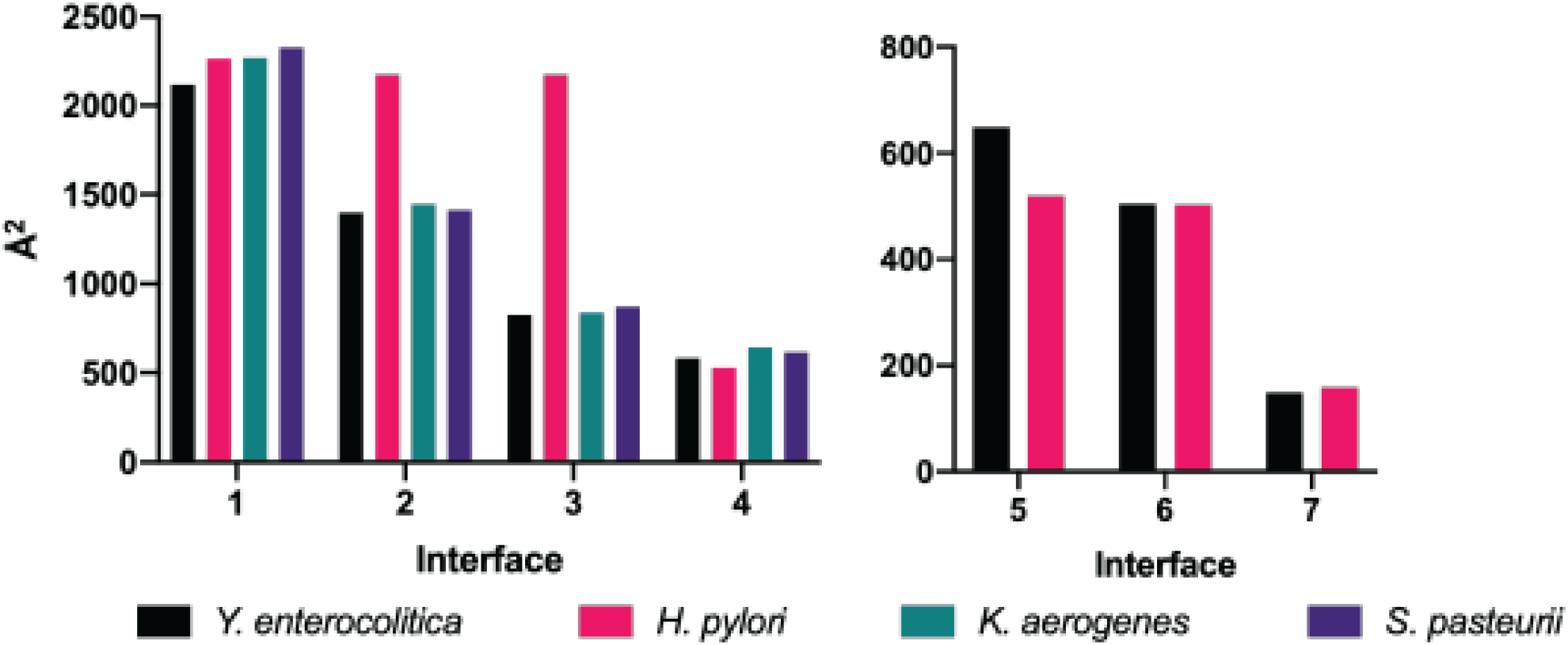
Area (Å^2^) of interfaces 1-7 (numbering as in Figure 4) plotted by organism. Left: intra-trimer interfaces; right: inter-trimer interfaces. The intra-trimer areas of interfaces 2 and 3 in *H. pylori* are higher because they are part of the same chain.

**Supplementary Figure 8.**
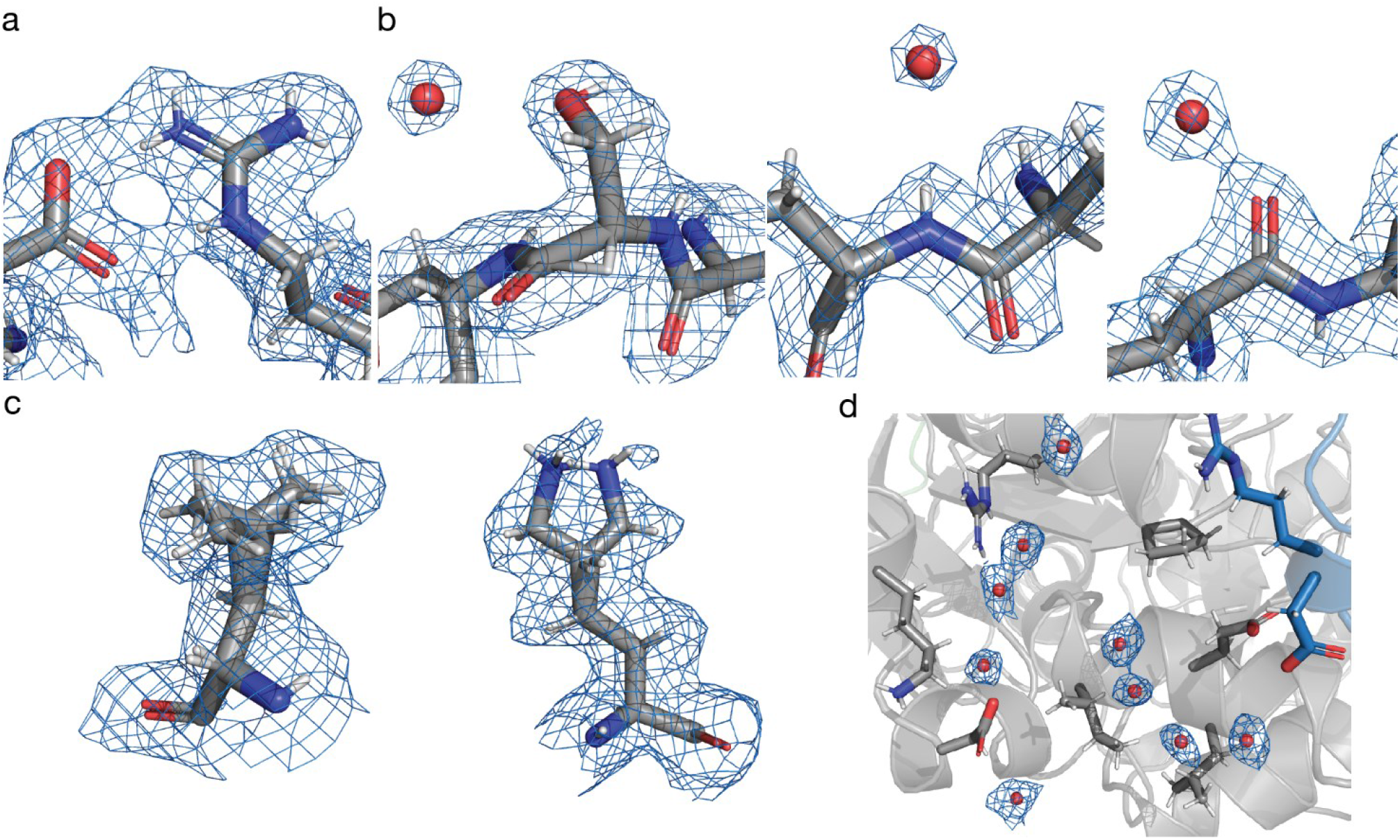
High-resolution structural features can be readily seen in the cryo-EM map. **a)** Salt bridge between D69 and R63. **b)** Side chain hydration of S17 and back bone hydration of the oxygen and the nitrogen, respectively. **c)** alternative side chain conformations of L406 and K2. d) network of waters between two ureC proteins (grey, blue).

**Supplementary Figure 9.**
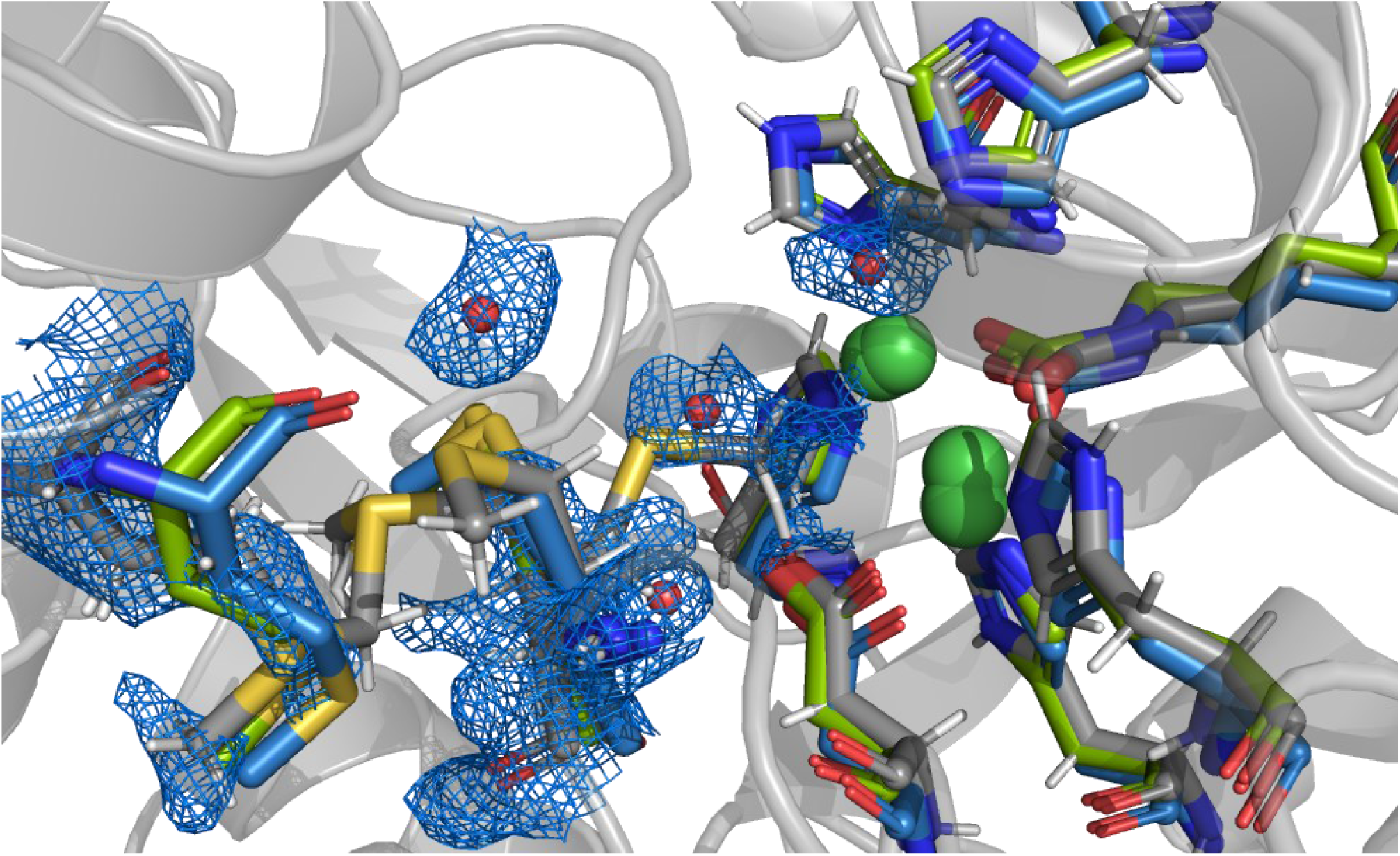
MET369 can adopt different conformations in the absence of the substrate or an inhibitor. *Y. enterocolitica* urease (grey), *S. pasteurii* urease (blue), *K. aerogenes* urease (green). One conformation could potentially reach the active site. There is no described function for this amino acid in the catalysis. The conformation close to the active site is only possible because the pocket is empty, which can be seen with an overlay of the active site from *K. aerogenes* urease (PDB: 1EJW) at 1.9 Å and *S. pasteurii* urease (PDB: 5OL4) at 1.28 Å resolution, respectively **(Figure 5d,e).**

### Supplementary Movies

**Supplementary Movie 1** Overview and feature highlights of the *Yersinia enterocolitica* urease cryo-EM structure.

**Supplementary Movie 2** Morphing between maps and models obtained from different sets of frames along the exposure.

### Supplementary Tables

**Supplementary Table 1.**
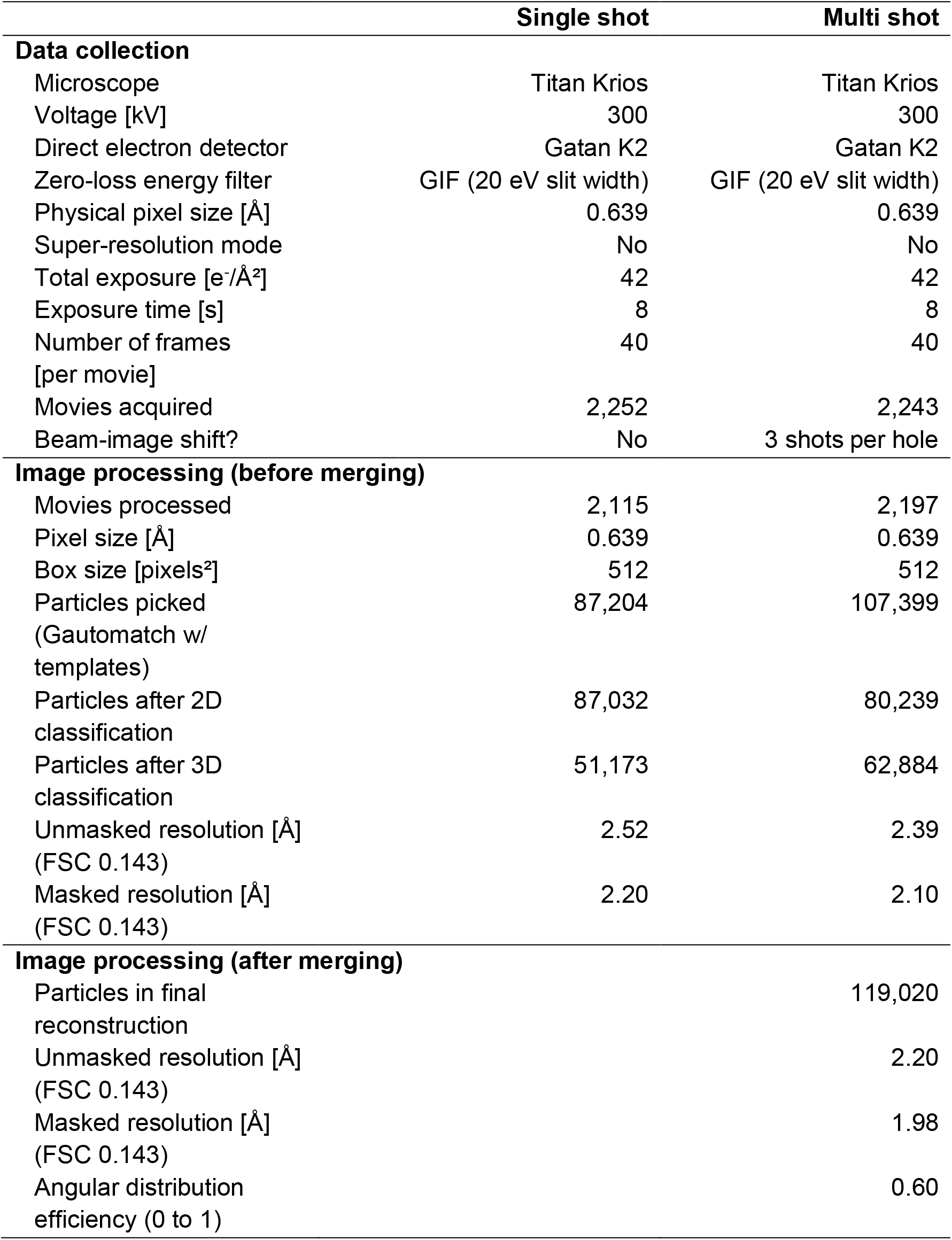
Cryo-EM data collection and image processing summary.

**Supplementary Table 2.**
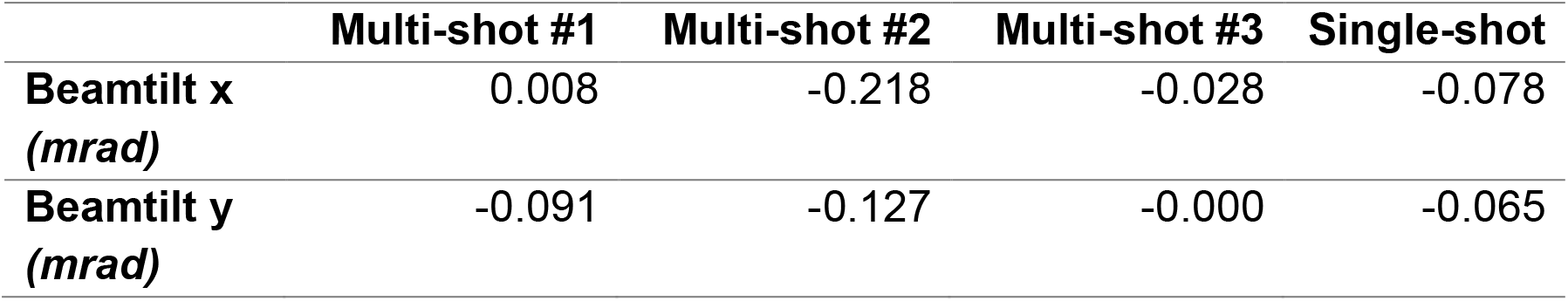
Beam tilt estimation after merging the two datasets.

|  | Multi-shot #1 | Multi-shot #2 | Multi-shot #3 | Single-shot |
| --- | --- | --- | --- | --- |
| Beamtilt x<br>(*mrad*) | 0.008 | -0.218 | -0.028 | -0.078 |
| Beamtilt y<br>(*mrad*) | -0.091 | -0.127 | -0.000 | -0.065 |

**Supplementary Table 3.** Bayesian polishing training results.

|  | **Multi**<br>**Shot**<br>**(before**<br>**merging)** | **Single**<br>**Shot**<br>**(before**<br>**merging)** | **Merged** | **Multi**<br>**Shot**<br>**#1** | **Multi**<br>**Shot**<br>**#2** | **Multi**<br>**Shot**<br>**#3** | **Single**<br>**shot** |
| --- | --- | --- | --- | --- | --- | --- | --- |
| **Number of particles**<br>**in training**<br>**(approximate)** | 5,000 | 5,000 | 10,000 | 10,000 | 10,000 | 10,000 | 10,000 |
| **Sigma for**<br>**Velocity (Å/dose)**<br>*(smaller = shorter*<br>*tracks)* | 0.531 | 0.819 | 0.779 | 0.810 | 0.693 | 0.636 | 0.681 |
| **Sigma for Divergence**<br>**(Å)**<br>*(higher = more*<br>*homogeneous tracks*<br>*across micrograph,*<br>*i.e. "rigid block")* | 3,600 | 10,455 | 10,620 | 10,260 | 7,350 | 5,220 | 8,355 |
| **Sigma for**<br>**Acceleration (Å/dose)**<br>*(smaller = straighter*<br>*tracks)* | 2.175 | 3.735 | 1.620 | 1.485 | 1.425 | 1.500 | 0.795 |

**Supplementary Table 4.** Sequence identity between different ureases and *Y. enterocolitica* urease.

| Organism | PDB code | Nr. Of Genes | Stoichiometry α-($\beta$)-($\gamma$) | ureA (%) | ureB (%) | ureC (%) |
| --- | --- | --- | --- | --- | --- | --- |
| *S. pasteurii* | 5O4L | 3 | 3-3-3 | 60.6 | 46.7 | 57.5 |
| *K. aerogenes* | 1EJW | 3 | 3-3-3 | 60.6 | 52.9 | 58.7 |
| *H. pylori* | 1E9Z | 2 | 12-12 | 52.0 | 50.4 | 57.6 |

**Supplementary Table 5.**
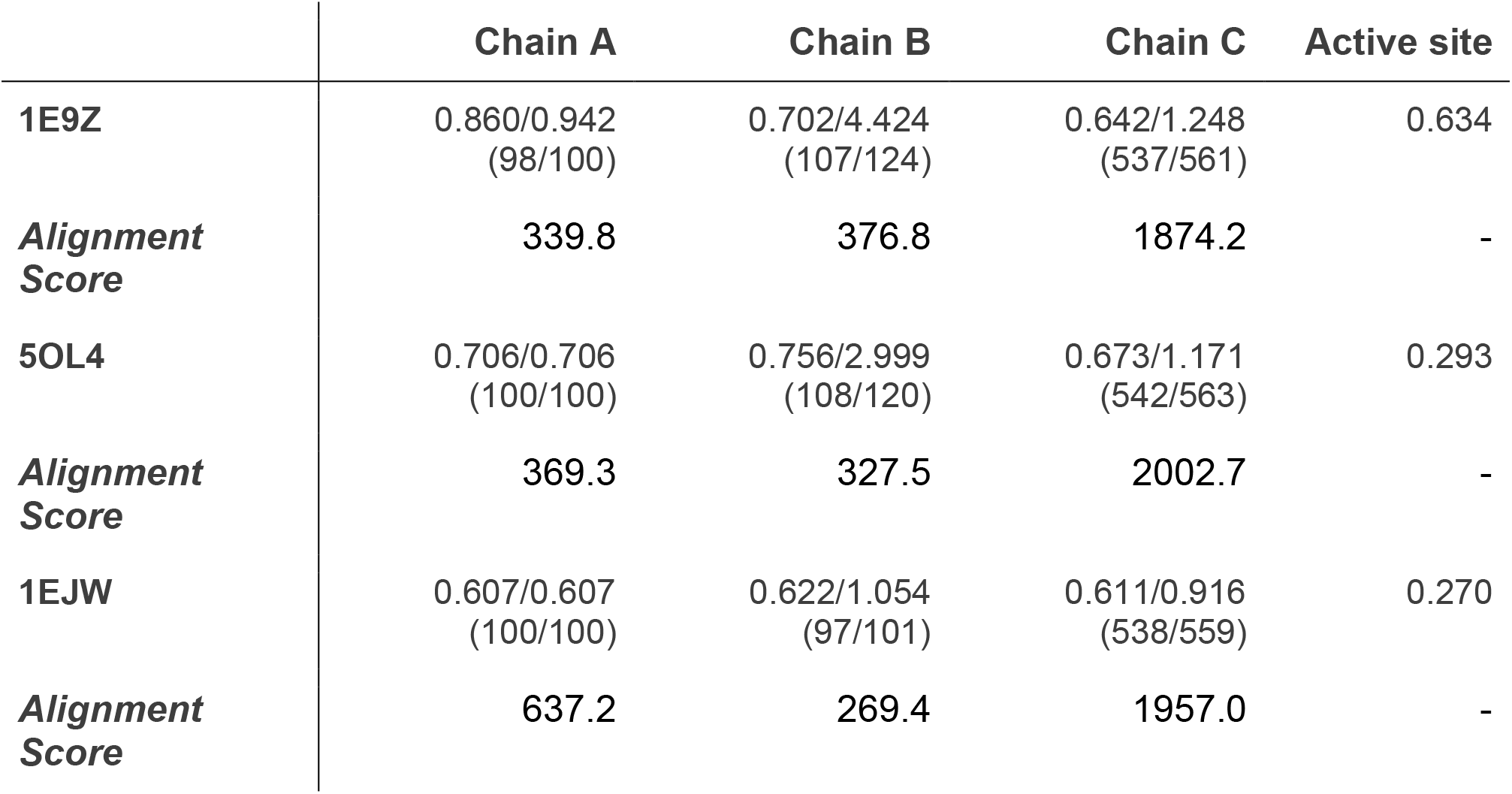
RMSD values between each chain of our *Y. enterocolitica* urease model versus structures deposited at the PDB. Values are given in Ångstroms. Values in parenthesis indicate the number of Cα atom pairs matched with and without pruning. The program matchmaker from UCSF Chimera was used to calculate sequence alignments and RMSD values between each chain in respective models.

|  | Chain A | Chain B | Chain C | Active site |
| --- | --- | --- | --- | --- |
| **1E9Z** | 0.860/0.942<br>(98/100) | 0.702/4.424<br>(107/124) | 0.642/1.248<br>(537/561) | 0.634 |
| **Alignment Score** | 339.8 | 376.8 | 1874.2 | - |
| **5OL4** | 0.706/0.706<br>(100/100) | 0.756/2.999<br>(108/120) | 0.673/1.171<br>(542/563) | 0.293 |
| **Alignment Score** | 369.3 | 327.5 | 2002.7 | - |
| **1EJW** | 0.607/0.607<br>(100/100) | 0.622/1.054<br>(97/101) | 0.611/0.916<br>(538/559) | 0.270 |
| **Alignment Score** | 637.2 | 269.4 | 1957.0 | - |

### Supplementary Note 1

**Supplementary Figure 10.**
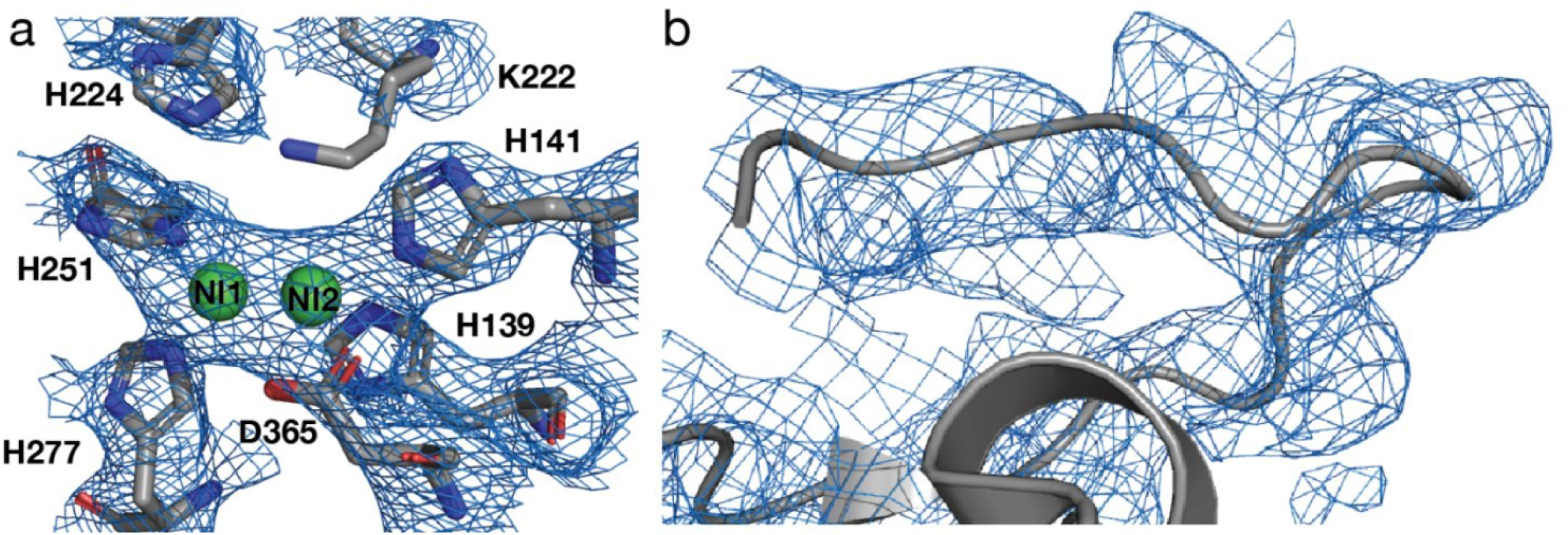
Low resolution crystal structure of *Y. enterocolitica* urease. Structure determination by X-ray crystallography yielded a 3.01 Å resolution density, which was sufficient to see the higher oligomeric state of *Y. enterocolitica* urease, but not for detailed analysis of the assembly mechanism and active site. Comparison of the model based on the high-resolution cryo-EM data *vs*. the previous crystal structure shows a few differences. The previous X-ray model did not include residue 100 of ureA and residues 31-33, 148-162 of ureB. Residues 148-162 form a helix, which was previously not observable. This helix sits right at the interface between three ureB proteins and constitutes the interaction between the trimers.

### Supplementary Methods

For X-ray crystallographic analysis of the *Y. enterocolitica* urease, *Y. enterocolitica* was differently purified than described in the **Methods** for cryo-EM. The protein was precipitated using 40-60% w/v ammonium sulfate, resuspended and dialyzed in 0.15M NaCl, 50mM Tris pH 8.0. It was further purified using a 45 ml self-packed DEAE FF XK26/20 (Sigma, DFF100) and a Superdex 200 10/300 GL column. The SEC was also used for buffer exchange to 20 mM HEPES, 100 mM NaCl pH 7. Urease crystals grew at 20°C at 10 mg/ml in 0.1M CHES pH 9.5; 50% (v/v) PEG 200. Urease crystals belonged to space group H32 with unit cell parameters of a= 157.2 Å, b = 157.2 Å and c= 774.6 Å, with four molecules per asymmetric unit. The structure was determined by molecular replacement with PHASER (McCoy *et al*., 2007) using the urease crystal structure from *Klebsiella aerogenes* (PDB: 1FWB) (Pearson *et al*., 1997). Model building and structure refinement were performed with Coot (Emsley & Cowtan, 2004) and Buster-TNT (Blanc *et al*., 2004). The atomic coordinates for this model have been deposited in the Protein Data Bank under the accession code 4Z42.

